# Dietary protein source alters gut microbiota composition and function

**DOI:** 10.1101/2024.04.04.588169

**Authors:** J. Alfredo Blakeley-Ruiz, Alexandria Bartlett, Arthur S. McMillan, Ayesha Awan, Molly Vanhoy Walsh, Alissa K. Meyerhoffer, Simina Vintila, Jessie L. Maier, Tanner Richie, Casey M. Theriot, Manuel Kleiner

**Author notes:** Please refer correspondence to J. Alfredo Blakeley-Ruiz or Manuel Kleiner. These authors contributed equally to this manuscript (J. Alfredo Blakeley-Ruiz, Alexandria Bartlett).

## Abstract

The source of protein in a person’s diet affects their total life expectancy. However, the mechanisms by which dietary protein sources differentially impact human health and life expectancy are poorly understood. Dietary choices have major impacts on the composition and function of the intestinal microbiota that ultimately modulate host health. This raises the possibility that health outcomes based on dietary protein sources might be driven by interactions between dietary protein and the gut microbiota. In this study, we determined the effects of seven different sources of dietary protein on the gut microbiota of mice using an integrated metagenomics-metaproteomics approach. The protein abundances measured by metaproteomics can provide microbial species abundances, and evidence for the molecular phenotype of microbiota members because measured proteins indicate the metabolic and physiological processes used by a microbial community. We showed that dietary protein source significantly altered the species composition and overall function of the gut microbiota. Different dietary protein sources led to changes in the abundance of microbial proteins involved in the degradation of amino acids and the degradation of glycosylations conjugated to dietary protein. In particular, brown rice and egg white protein increased the abundance of amino acid degrading enzymes. Egg white protein increased the abundance of bacteria and proteins usually associated with the degradation of the intestinal mucus barrier. These results show that dietary protein sources can change the gut microbiota’s metabolism, which could have major implications in the context of gut microbiota mediated diseases.

## INTRODUCTION

Source of dietary protein has a major impact on human health. People who consume high amounts of animal protein have higher mortality rates than those who consume mostly plant-based protein [1, 2]. Egg protein and red meat protein in particular have been shown to lead to increased mortality rates among humans [3] and a diet high in red meat protein has been shown to increase inflammation in a mouse model of colitis [4]. Replacing animal protein sources with plant protein sources in humans reduces mortality rates [3]. Currently, we have a limited understanding of the underlying causes, but the gut microbiota has been implicated as potentially having a major role in the differential health impacts of different dietary protein sources [5, 6]. Diet has been shown to change the gut microbiota’s composition and function in ways that can be detrimental or beneficial to health in both mice and humans[7–10]. For example, protein fermentation by the gut microbiota generates a number of toxins including ammonia, putrescine, and hydrogen sulfide [6, 11], whereas fermentation of fiber and certain amino acids produces anti-inflammatory short-chain fatty acids [12]. Previous studies in mice demonstrate that the amount of protein can have a greater impact on the gut microbiota’s composition than other macronutrients [13], and that source of dietary protein alters gut microbiota composition[14]. There is, however, limited data showing the mechanisms by which individual sources of dietary protein affect the gut microbiota’s composition and function, which could mediate the consumption and production of compounds beneficial or detrimental to the host.

Metaproteomics represents a powerful tool for characterizing the mechanisms underlying dietary effects on gut microbiota [7, 9]. Metaproteomics is defined as the large-scale characterization of the proteins present in a microbiome [15]. Protein abundances measured by metaproteomics simultaneously provide microbial species abundances [16], and evidence for the metabolic and physiological phenotype of microbiota members [17–19]. Metaproteomes are usually measured using a shotgun proteomics approach where proteins extracted from a sample are digested into peptides, separated by liquid chromatography, and measured on a mass spectrometer [20]. Proteins are then identified and quantified using a database search algorithm, which matches the measured peptides to a database of protein sequences [21]. Due to the heterogeneous nature of complex microbial communities it is usually best to construct the protein database using gene predictions from metagenomes measured from the same samples [21]. When metaproteomics is coupled to a genome-resolved metagenomic database it is possible to evaluate strain and species level function even if the microbes have not been previously characterized [22, 23]. We call this approach integrated metagenomics-metaproteomics.

We used an integrated metagenomic-metaproteomic approach to investigate the effects of dietary protein source on gut microbiota composition and function. We hypothesized that dietary protein source affects the abundance of amino acid metabolizing enzymes from the gut microbiota, altering the abundance of pathways involved in the production of toxins detrimental to host health. We found that the source of dietary protein not only alters the abundance of amino acid degrading enzymes, but has an even greater impact on the abundance of glycan degrading proteins among other functions, indicating that dietary protein sources can have wide ranging effects on the gut microbiota.

## MATERIALS AND METHODS

### Experimental overview, description of diets, animal housing and preliminary results

We used twelve C57BL/6J mice in two groups (six males, six females, Jackson Labs, Bar Harbor) aged three months at the beginning of the study. The males and females originated from different mouse rooms at the Jackson Labs and thus were expected to have different background microbiomes. Mice from both groups were housed in two separate cages (3 mice/cage) with a 12 h light/dark cycle. We autoclaved bedding, performed all cage changes in a laminar flow hood and maintained an average temperature of 70°F and 35% humidity. We conducted our animal experiments in the Laboratory Animal Facilities at the NCSU CVM campus (Association for the Assessment and Accreditation of Laboratory Animal Care accredited), which are managed by the NCSU Laboratory Animal Resources. Animals assessed as moribund were humanely euthanized via CO_2_ asphyxiation. NC State’s Institutional Animal Care and Use Committee approved all experimental activity (Protocol # 18-034-B).

We fed the mice a series of nine fully defined diets (Fig. 1a, Supplementary table 1) formulated and purchased from Envigo Teklad (inotiv, Madison, WI, USA) based on the diets used a previous study. We worked with Envigo to formulate 20% or 40% protein (by weight) diets, which required slight variations in the amount of protein source added for each protein source. The torula yeast protein source has large amounts of heavily glycosylated mannoprotein which resulted in a higher amount of the yeast protein source being added to achieve a 20% protein content (Supplementary table 1). We chose 20% and 40% as the protein amounts for this study partially due to the results of a previous study which showed that most of the effects of amount of protein on the gut microbiota happened between 6% and 20% and that there was not much of an effect from 20% to 40% [24]. Because the goal of the study was to investigate the effects of source of dietary protein, we selected 20% protein as our main dietary protein amount to avoid any unwanted microbiota effects due to small variations in amount of protein. We included two additional diets with 40% as the amount of both to serve as an internal control by providing the dietary protein source again and also to show that the increase from 20% to 40% would not have too much of an effect on the microbiota. In order of feeding, the diets were 20% soy protein, 20% casein protein, 20% brown rice protein, 40% soy protein, 20% yeast protein, 40% casein protein, 20% pea protein, 20% egg white protein, and 20% chicken bone protein. To control for the succession effects of the serial dietary intervention and the age of the mice, we fed the mice the 20% soy diet or the 20% casein diet as an additional control at the end of the diet series. In order of feeding, the diets were 20% soy protein, 20% casein protein, 20% brown rice protein, 40% soy protein, 20% yeast protein, 40% casein protein, 20% pea protein, 20% egg white protein, and 20% chicken bone protein. To control for the succession effects of the serial dietary intervention and the age of the mice, we fed the mice the 20% soy diet or the 20% casein diet as an additional control at the end of the diet series. The chicken bone diet caused the mice to lose weight, so we discontinued the diet after 3 days and the mice consumed a standard chow diet for the rest of that week. No fecal samples were collected for the chicken bone diet. All defined diets were sterilized by ɣ-irradiation and mice were provided sterile water (Gibco). After feeding each diet for 7 days, we collected fecal samples, prior to replacing food with the next diet. We collected samples in NAP preservation solution at a 1:10 sample weight to solution ratio, and roughly homogenized the sample with a disposable pestle prior to freezing at -80°C [25]. We had to sacrifice one mouse during the second diet (20% casein) so no additional samples were collected. We also were unable to collect a sample from one of the mice during the brown rice and egg white diets so only 10 samples were collected for those diets. We analyzed samples from all mice and diets using an integrated metagenomic-metaproteomic approach [7, 23] (Fig. 1b).

**Figure 1:**
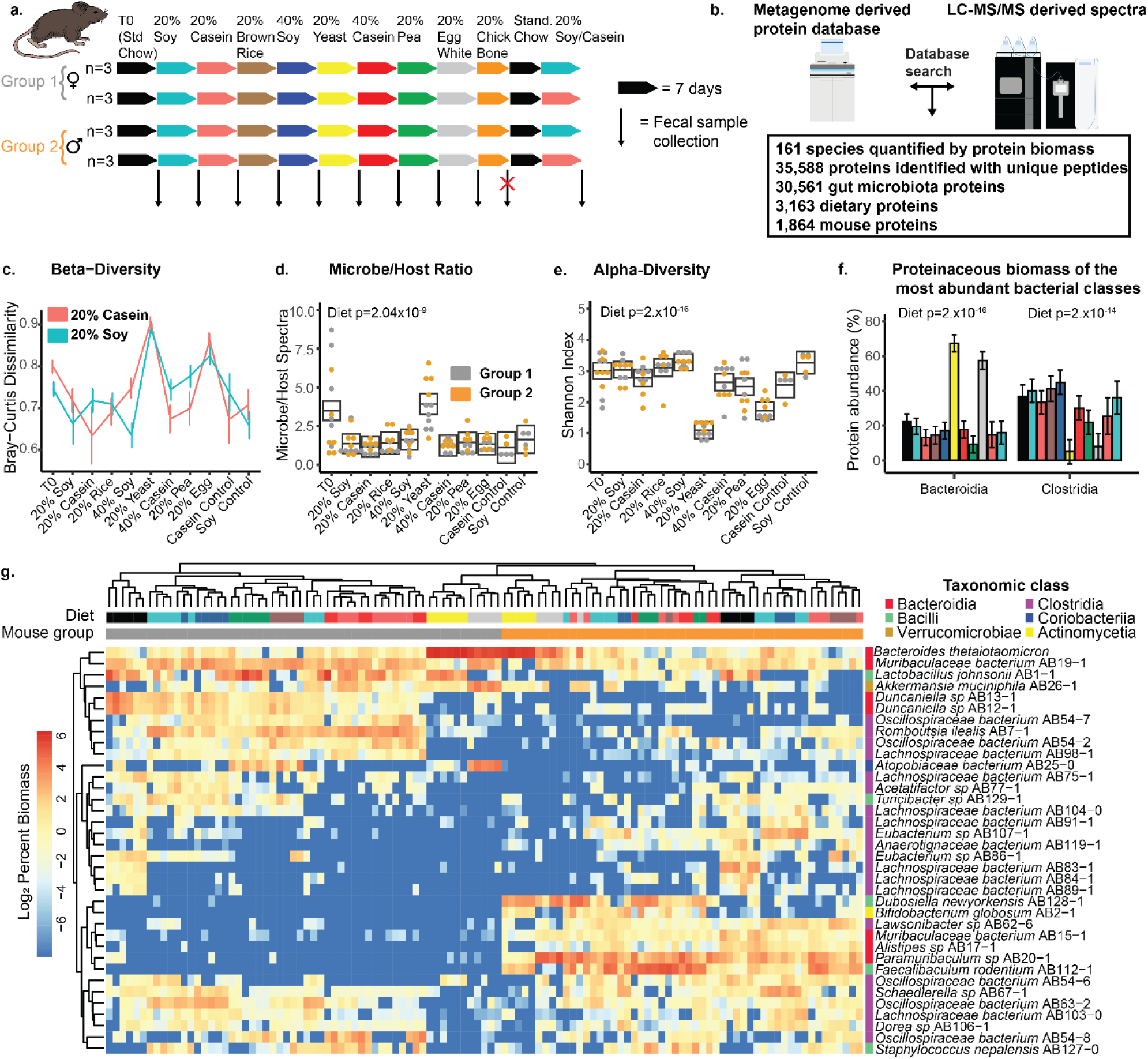
Source of dietary protein alters the gut microbiota’s composition. (a) Diagram showing the experimental design, with number of cages, and order of the diets fed. Colors depicting the diets are used throughout the manuscript. Each row of arrows represents one cage. We collected 10-12 samples for each experimental diet and 5-6 samples for each control diet. (b) A diagram illustrating the integrated metagenomic-metaproteomics method used to analyze the samples along with raw metrics: quantifiable species and number of proteins. (c) Bray-Curtis dissimilarity between the initial 20% soy diet (teal) or 20% casein diet (red) and all other diets. Error bars reflect 95% confidence intervals for all line graphs as calculated by the Rmisc package in R, but the PERMANOVA values from Table 1 and Extended Data Table 5 are the true tests of significance. (d) Ratio of spectra assigned to microbes versus the host; boxes represent 95% confidence intervals calculated on a linear mixed effect model (Extended Data Table 2). (e) Shannon diversity index of the gut microbiota across all diets; boxes represent 95% confidence intervals calculated on a linear mixed effect model; boxes that do not overlap are significantly different (Extended Data Table 2). (f) Abundances of the two most abundant bacterial classes *Bacteroidia* and *Clostridia* based on summed protein abundance. Colors represent diets and the legend is (a). Error bars are 95% confidence intervals calculated using a linear-mixed effects model and error bars that do not overlap represent significant differences (Extended Data Table 2). (g) A hierarchically clustered (ward.D2 algorithm on euclidean distances) heatmap depicting the clustering by species group abundance of the 36 most abundant species in the study. Species were considered abundant if they had at least 5% of the microbial biomass in at least one sample.

**Table 1:** PERMANOVA analysis of the factors that explain the variance proteinaceous biomass gut microbiota species. Table of proteinacieous biomass of species was analysed using the adonis2 function in the vegan package in R. Similarity matrix was calculated using Bray-Curtis and the factors were protein source, amount of protein, mouse group, and age in weeks. The strata function was used to account for repeated measures. *R*^2^ describes how much of the variance is explained by the factor, whereas F determines if the difference is significant.

| Factor | df | SumOfSqs | $R^2$ | F | P |
| --- | --- | --- | --- | --- | --- |
| Protein source | 7 | 12.087 | 0.39 | 14.95 | 0.001 |
| Amount of protein | 2 | 0.565 | 0.01 | 2.2 | 0.003 |
| Mouse group | 1 | 7.118 | 0.21 | 56.9188 | 0.001 |
| Mouse age | 1 | 0.317 | 0.0095 | 2.533 | 0.017 |

### Metagenomic DNA sequencing

To create a database for metaproteomic analysis, we pooled fecal samples from each cage to create four cage specific metagenomes. The metagenomes were only used to generate a protein sequence database for metaproteomics, and all results presented in the manuscript are derived from protein quantities acquired by metaproteomic measurements. We gathered one fecal sample from each cage for four different diets (20% rice, 40% soy, 20% yeast, 40% casein) for a total of 16 samples.To remove the preservation solution from the samples, we added 5 mL of 1X Phosphate Buffered Saline solution (VWR) to the samples and centrifuged them (17,000 x g, 5 min) to pellet solids and bacterial cells in suspension. We removed the preservation solution and resuspended the fecal pellets in 1 mL of InhibitEX Buffer in Matrix E (MP Biomedicals) bead beating tubes. We beat the samples at 3.1 m/s for 3 cycles with 1 minute of ice cooling between each cycle using a Bead Ruptor Elite 24 (Omni International). We isolated DNA from the resulting lysate using the Qiagen QIAamp Fast DNA stool mini kit (cat. No. 51604) [26]. Samples were extracted individually and pooled by cage with each sample contributing a total of 200 ng of DNA.

We submitted genomic DNA to the North Carolina State Genomic Sciences Laboratory (Raleigh, NC, USA) for Illumina NGS library construction (TruSeq Nano Library kit) and sequencing (NovaSeq 6000 sequencer). We obtained between 51,152,549 and 74,618,259 paired-end reads for each of the 4 samples.

### Metagenomic assembly and protein database construction

We assembled raw reads using a genome resolved metagenomics approach. We removed PhiX174 (NCBI GenBank accession CP004084.1) and mouse genome (mm10) contaminating sequences using BBSplit and removed adapters using BBDuk (BBMap, Version 38.06), parameters: mink = 6, minlength =20 [27]. We assembled decontaminated reads individually using MetaSPAdes (v3.12.0) -k 33,55,99 [28] and co-assembled them using MEGAHIT (v1.2.4) --kmin 31 --k-step 10 [29]. We mapped reads from all four samples to all five assemblies using bbmap, and binned the contigs using MetaBAT (v2.12.1) [30]. We assessed the quality of the bins using CheckM (v1.1.3) [31] and automatically accepted medium quality bins with a completion score greater than 50% and a contamination score of less than 10% [32]. Because the purpose of metagenomics in our study was to generate a comprehensive protein sequence database and to assign proteins to species, we further accepted bins that were greater than >30% complete and <5% contaminated. We do not think that genomes of lower than quality than medium, should be contributed to major public databases like those hosted by NCBI and ENA so we have made these MAGs available in Dryad instead (see data availability section). We clustered the bins into species groups by 95% ANI using dRep (v2.6.2) [33, 34] and assigned taxonomy using GTDB-Tk (v1.3.0, ref r95) [35].

We assembled the protein database by combining gene annotations from the metagenome with mouse and dietary protein databases [21]. For the metagenome, we annotated the assemblies prior to binning and then for each bin individually using PROKKA (Version 1.14.6)[36]. If the contig was binned, we compiled the annotations from the bins. We then used CD-HIT-2D (Version 4.7), with a 90% identity cutoff, to compare the genes from the unbinned PROKKA output to the binned gene annotation [37]. If a gene was not present in a bin we added it to the database as an unbinned sequence. Once we compiled the microbial protein database we assigned each protein sequence a species code if it was species specific or an ambiguous, low-quality, or unbinned code if it was assigned to more than one species group, belonged to a low-quality bin, or was not present in a bin. In addition to the microbial sequences, we added a *Mus musculus* proteome (UP000000589, Downloaded 19Feb20), and the relevant dietary protein database for each sample: *Glycine max* (UP000008827, Downloaded 19Feb20), *Bos taurus* (UP000009136, Downloaded 19Feb20), *Cyberlindnera jadinii* (UP000094389, Downloaded 25May20), *Oryza sativa* (UP000059680, Downloaded 25May20) and *Gallus gallus* (UP000000539, Downloaded 25May20). Due to the lack of a reference proteome for the yellow pea diet, we created a custom pea reference with all available UniProtKB [38] protein sequences for *Pisum sativum* (Taxon ID: 388 Downloaded 25Apr20) and the reference proteome of *Cajanus cajan* (UP000075243, Downloaded 25May20). For T0 samples taken when mice were fed a standard chow diet, we added proteomes from the protein sources likely to be in the diet based on the ingredient list (corn UP000007305, fish UP000000437, soy UP000008827, wheat UP000019116, Downloaded 19Feb20). We clustered the mouse and diet reference proteomes individually at a 95% identity threshold. We only searched samples against their respective dietary database. The databases ranged in size from 597,215 to 809,468 protein sequences depending on the dietary components, each database had the same number of microbial and host proteins.

In order to identify all sequences from the species *Bacteroides thetaiotaomicron* and *Lactococcus lactis* we downloaded all the sequences matching these species from UniProt [38]. We then used diamond BLASTp to identify all sequences in the protein database that matched with 95% identity or greater. The species code for these sequences was changed to BT or LAC if they were found to be *B. thetaiotaomicron* or *L. lactis* respectively.

### Metaproteomic sample processing

We extracted protein using a modified FASP protocol[39]. We pelleted fecal samples by centrifugation (21,000 x g, 5 min) and removed the preservation solution. We suspended dietary and fecal pellets in SDT lysis buffer [4% (w/v) SDS, 100 mM Tris-HCl pH 7.6, 0.1 M DTT] in Lysing Matrix E tubes (MP Biomedicals) and bead beat the samples (5 cycles of 45 s at 6.45 m/s, 1 min between cycles). After bead beating we heated the lysates to 95°C for 10 minutes. We mixed 60 µL of the resulting lysates with 400 µL of UA solution (8 M urea in 0.1 M Tris/HCl pH 8.5), loaded the sample onto a 10 kDa 500 µL filter unit (VWR International) and centrifuged at 14,000 x g for 30 minutes. We repeated this step up to three times to reach filter capacity. After loading, we added another 200 µL of UA buffer and centrifuged at 14,000 x g for another 40 minutes. We added 100 µL of IAA solution (0.05 M iodoacetamide in UA solution) to the filter and incubated at 22°C for 20 minutes. We removed IAA by centrifuging the filter at 14,000 x g for 20 minutes. We then washed the filter 3 times by adding 100 uL of UA buffer and centrifuging at 14,000 x g for 20 minutes. We then washed 3 more times by adding 100 uL of ABC buffer (50 mM Ammonium Bicarbonate) and centrifuging at 14,000 x g for 20 minutes. To digest the isolated protein, we added 0.95 µg of MS grade trypsin (Thermo Scientific Pierce, Rockford, IL, USA) mixed in 40 µL of ABC to each filter and incubated at 37°C for 16 hours. We eluted the peptides by centrifugation at 14,000 x g for 20 minutes. We eluted again with 50 uL of 0.5 M NaCL and centrifuged at 14,000 x g for another 20 minutes. We quantified the abundance of the peptides using the Pierce Micro BCA assay (Thermo Scientific Pierce, Rockford, IL, USA) following the manufacturer’s instructions.

We analyzed the samples by 1D-LC-MS/MS. Samples were run in randomized block design. For each run, we loaded 600 ng of peptides onto a 5 mm, 300 µm ID C18 Acclaim® PepMap100 pre-column (Thermo Fisher Scientific) using an UltiMate 3000 RSLCnano Liquid Chromatograph (Thermo Fisher Scientific) and desalted on the pre-column. After desalting, the pre-column was switched in line with a 75 cm x 75 µm analytical EASY-Spray column packed with PepMap RSLC C18, 2 µm material (Thermo Fisher Scientific), which was heated to 60 °C. The analytical column was connected via an Easy-Spray source to a Q Exactive HF Hybrid Quadrupole-Orbitrap mass spectrometer. Peptides were separated using a 140 minute reverse phase gradient [24]. We acquired spectra using the following parameters: m/z 445.12003 lock mass, normalized collision energy equal to 24, 25 s dynamic exclusion, and exclusion of ions of +1 charge state. Full MS scans were acquired for 380 to 1600 m/z at a resolution of 60,000 and a max IT time of 200 ms. Data-dependent MS^2^ spectra for the 15 most abundant ions were acquired at a resolution of 15,000 and max IT time of 100 ms.

### Metaproteomic data processing

We searched raw MS spectra against the diet specific protein databases using the run calibration, SEQUEST HT and percolator nodes in Proteome Discoverer 2.3 (Thermo Fisher Scientific). We used the following setting for search: trypsin (full), 2 missed cleavages, 10 ppm precursor mass tolerance, 0.1 Da fragment mass tolerance. We included the following dynamic modifications: oxidation on M (+15.995 Da), deamidation on N,Q,R (0.984 Da) and acetyl on the protein N terminus (+42.011 Da). We also included the static modification carbamidomethyl on C (+57.021 Da). We filtered identified peptides and proteins at a false discovery rate (FDR) of 5%. Additionally, we only included proteins that had at least one protein unique peptide identified. Proteins were quantified by peptide spectral match (PSM) count (spectral counting).

### Statistical analysis and visualization

Whenever possible we tested significance of changes in abundance by applying an ANOVA on a linear mixed effects model with the interacting fixed effects being mouse group and diet, and the random effect being the individual mouse (lme4 version 4.3.1) [40]. For multiple comparisons we calculated 95% confidence intervals for each diet using the emmeans R package (version 1.8.8) [41]. The exceptions were PERMANOVA analysis for testing significance of microbiota compositional changes (Table 1; Extended Data Table 5) and Welch‘s t-tests to compare differences between yeast and egg white protein diets (Extended Data Table 17, 18 and 20). For each analysis, we controlled for multiple-hypothesis testing by converting *P* values to FDR-level based q-values, unless all *P* values in the analysis were below 0.05 using Benjamini-Hochberg procedure [42, 43]. By definition, if all the *P* values are less than 0.05 than the FDR is less than 0.05. Visualizations were produced using ggplot2 (version 3.4.3) [44], pheatmap (version 1.0.12) [45], RawGraphs [46], Microsoft Excel and Adobe Illustrator. All boxes and error bars represent 95% confidence intervals. Boxes or error bars that do not overlap denote significance. If no error bars are present then significance is denoted by letters or asterisk. In the case of beta-diversity analysis the error bars are 95% confidence intervals (Fig. 1c), but significance was tested separately by PERMANOVA.

### Compositional profiling of gut microbiota

We calculated the abundances of gut microbes using proteinaceous biomass [16]. Briefly, we filtered for proteins with at least 2 protein unique peptides and summed their spectra into their assigned taxonomy: microbial species, mouse, diet, ambiguous, low quality bins, unbinned bacteria. We calculated the microbe to host ratio by summing the spectral count assigned to microbial species, multiple microbial species (ambiguous), low quality bins and unbinned bacteria proteins and dividing them by the number of spectral counts assigned to mouse proteins. We considered a microbial species quantifiable if we could identify at least one protein with 2 protein unique peptides unambiguously assigned to the species. If a protein matched to more than one species, it was not included in the quantification. We showed previously that these filtering and inclusion/exclusion criteria allow for accurate biomass estimates [16].We calculated per sample species richness by simply counting the number of quantifiable species per sample. We calculated alpha (Shannon Diversity Index) and beta diversity (Bray-Curtis) metrics using the vegan (version 2.6-4) package in R (version 4.3.1) [47, 48] on a table of the quantifiable microbial species (statistics as described above). We also evaluated gut microbiota composition using principal component analysis and hierarchical clustering. For principal component analysis we normalized the quantified species using centered-log ratio transformation and calculated principal components using the prcomp function in base R on all the mice and separately on each mouse group. Principal components were rendered using the ggplot2 (version 3.4.3) package in R [44]. For hierarchical clustering, we focused on the species that were most abundant, representing at least 5% of the microbial species biomass in at least one sample. We calculated the percent biomass for all the species and then extracted the species that fit the abundant species criteria. We calculated the individual significance of each abundant species using linear mixed effects models as described above. We hierarchically clustered log transformed values of these species using the R package pheatmap (version 1.0.12), using the ward.D2 algorithm and euclidean distances [45]. To compare broad taxonomic changes at the class level, for all quantifiable species, we summed the abundance of the assigned class by GTDB-Tk.

### Functional profiling of gut microbiota

For analyses of functional categories at the level of the whole microbiota we calculated the normalized spectral abundance factor (NSAF%) for each protein, which provides the relative abundance for each protein as a percentage of the summed abundance of microbiota proteins [49]. We annotated functions for all microbial proteins in our database using EggNOG-mapper [50], MANTIS [51], and Microbe Annotator [52]. We assigned glycoside hydrolase protein family identifiers from the CAZy database using dbCAN2 [53, 54]. We manually curated these annotations by searching a subset of these proteins against the Swiss-Prot [38] and InterPro [55] databases between February 2023 and June 2023. If the Swiss-Prot or InterPro annotations matched the automated tool annotations we extrapolated the assigned protein name to all proteins with the same automated annotation. Alternatively, if the annotations from the automated tools were in agreement, we consolidated the annotation into a consensus annotation. We then assigned broad functional categories, detailed functional categories, and specific names to each validated protein set. To evaluate functional changes due to diet, we summed all microbiota proteins assigned to a broad or detailed functional category, or enzyme name and applied a linear mixed-effects model to each function.

### In vivo proteomic analysis of B. thetaiotaomicron

To analyze the *B. thetaiotaomicron* proteome, we calculated the orgNSAF by extracting all proteins assigned to the species *B. thetaiotaomicron* detected in the metaproteomes, and then calculating NSAF% [17]. We then compared abundances of *B. thetaiotaomicron* proteins detected in the yeast and egg protein diets using the Welch’s t-test in the Perseus software (version 1.6.14.0) [56]. To visualize polysaccharide utilization loci (PULs) we mapped the reads from one of our metagenomic samples to all the contigs that were assigned *B. thetaiotaomicron* proteins using BBSplit (BBMap, Version 38.06). We then assembled all the mapped reads using metaSPAdes. The genes in this newly assembled genome overlapped exactly with the previous set of identified *B. thetaiotaomicron* genes, and this *B. thetaiotaomicron* genome was uploaded to the RAST server for further analysis [57]. PULs were detected in the metaproteome by identifying proteins labeled SusC, SusD, or TonB. The rest of the PUL was identified by visualizing the gene neighborhood in RAST. The identified genes were then cross referenced against PULDB to assign literature described PUL numbers [58].

### In vitro growth and proteomics of B. thetaiotaomicron

We cultured *B. thetaiotaomicron* VPI-5482 in two biological replicates and at least 4 technical replicates using a defined *Bacteroides* medium [59]. *B. thetaiotaomicron* cultures were grown statically at 37°C in a Coy anaerobic chamber (2.5 % H_2_ /10 % CO_2_ /88.5 % N_2_) in minimal medium (100 mM KH_2_PO_4_, 8.5 mM [NH_2_]_4_SO_4_, 15 mM NaCl, 5.8 μM vitamin K_3_, 1.44 μM FeSO_4_⋅7H_2_O, 1 mM MgCl_2_, 1.9 μM hematin, 0.2 mM L-histidine, 3.69 nM vitamin B_12_, 208 μM L-cysteine, and 7.2 μM CaCl_2_⋅2H_2_O). The four dietary protein sources: soy protein (CA.160480), yeast protein (CA.40115), casein protein (CA.160030), and egg white protein (CA.160230), were purchased from Envigoand were the same as the protein sources used in the corresponding diets. Porcine muc2 mucin (Sigma CA. M2378) was also tested alongside controls of glucose and no carbon source. To aid in suspension in aqueous media, we pre-prepared the proteins in 200 mM NaOH water at 37°C for four days; the glucose control was also dissolved in 200 mM NaOH water. We then added the protein or glucose solution to the pre-prepared media at 0.5% (wt/v). Cultures were grown overnight in minimal media supplemented with 0.5% (wt/v) glucose before being washed and inoculated into experimental conditions at 0.01 OD and incubated at 37°C anaerobically with shaking every hour. Colony forming units (CFUs) per mL of culture were enumerated by drip plating at 0 and 24 hr post inoculation. Solid media for *B. thetaiotaomicron* was Brain-Heart Infusion agar (Difco CA. 241830) supplemented with 10% Horse Blood (LAMPIRE CA. 7233401) (BHI-HB).

To obtain samples for proteomics, we repeated the experiment for the glucose, yeast, egg white, mucin and soy media. After 8 hours, CFUs were enumerated to confirm growth. We pelleted cells by centrifuging at 4,000 g for 10 minutes. We then extracted the supernatant and froze the pellets at -80°C. Protein was extracted by the same FASP protocol described above but with two differences. We lysed pellets by adding 120 uL of SDT buffer and then heating at 95°C. We used PES 10kDa filters (MilliporeSigma). We also used a similar Mass Spectrometry procedure, except the samples were run on an Exploris 480 mass spectrometer (Thermo Fisher Scientific) and 1 ug of peptide were analyzed for each sample. We searched raw MS spectra using the same Proteome Discoverer 2.3 workflow using the *B. thetaiotaomicron* proteome downloaded from UniProt (UP000001414 downloaded January 9, 2024) as the protein sequence database. We cross referenced PULs between the metaproteome and the *in vitro* proteome to compare them.

## RESULTS

### Source of dietary protein alters gut microbiota composition

Shotgun sequencing of the fecal samples using a genome-resolved metagenomics pipeline [33, 60] resulted in 454 metagenome-assembled genomes (MAGs) organized into 180 species groups. We used high-resolution mass spectrometry based metaproteomics to identify and quantify proteins in each sample using a protein sequence database derived from the metagenome and augmented with mouse and diet protein sequences [20, 21]. In total, we identified 35,588 proteins, each distinguished as microbial, host, or dietary proteins (Extended Data Table 1). We quantified the proteinaceous biomass for each species using metaproteomics data and obtained measurements for 161 distinct species (Extended Data Table 3) [16]. PERMANOVA analysis on the 161 quantified species revealed that protein source and mouse group best explained the variance in gut microbiota composition with *R*^2^ values of 0.39 and 0.29 respectively, whereas amount of protein in the diet and age of the mouse had little effect (*R*^2^ values 0.01 or less (Table 1). Pairwise comparisons using Bray-Curtis showed that the source of dietary protein significantly changed the composition of the gut microbiota in 43 out of 49 comparisons (FDR controlled PERMANOVA q < 0.01; Extended Data Table 5), and that the yeast and egg white diets had the most dissimilar gut microbial communities(Fig. 1c). In all cases our internal controls 40% soy and casein diets in the middle of the study, and the 20% soy and casein control feedings at the end of the study had the least dissimilar Bray-Curtis values when compared to the initial 20% soy or 20% casein diets respectively, which helped to validate the initial rationale described in the methods. We also assessed the ratio of microbial to host proteins as proxy for bacteria load and the alpha diversity (within sample diversity) using the Shannon Diversity index. We found that bacterial load significantly increased in the yeast diet (Fig. 1d) and that within sample diversity significantly decreased in the yeast and egg white diets (Fig. 1e).

The large differences in microbial composition in the egg white and yeast protein diets were driven by a decrease in the abundances of species from the class *Clostridia* in favor of species from the class *Bacteroidia* (Fig. 1f). Because we observed fewer species in the class *Bacteroidia* overall, it makes sense that a drop in *Clostridia* in favor of *Bacteroidia* would result in a lower alpha diversity in the yeast and egg white diets (Fig. 1g; Extended Data Table 3).

We analyzed the most abundant microbial species (>5% of the microbial protein biomass in at least one sample) and hierarchically clustered them by abundance across the different dietary protein sources/groups (Fig. 1g). This revealed three major clusters separating most samples by mouse group with the exception of the yeast and egg white diets which together formed a separate cluster that internally showed separation by mouse group. The T0 samples fell into the major mouse group clusters which indicates that the two mouse groups had distinct gut microbial communities at the start of the experiment. Within the mouse group clusters the abundance of gut microbes clustered by source of dietary protein, which was also observed in principal component analysis (Supplementary Fig. 1, Supplementary Results Section A). In the yeast diet, *B. thetaiotaomicron* dominates gut microbiota composition regardless of mouse group. *B. thetaiotaomicron* abundance also increased in response to the egg white diet. There were additional species specific to each mouse group that also increased in abundance in response to the egg white diet (Fig. 1g; Supplementary Fig. 2). In group 1, these species were *Akkermansia muciniphila* and *Atopobiaceae* bacterium AB25-9, but in group 2 these species were *Paramuribaculum* sp. and *Dubosiella newyorkensis.* Both *A. muciniphila* [61] and *Paramuribaculum* sp. [62] have been reported to forage on intestinal mucin and *B. thetaiotaomicron* has been shown to switch towards mucin foraging when fed a low-fiber diet [63]. These results show that the source of dietary protein changes the gut microbiota’s composition and suggests that an egg white diet could promote mucin-foraging bacteria.

### Source of dietary protein alters gut microbiota function

We used the normalized abundances of gut microbiota proteins as a measure of the investment into metabolic and physiological functions [17, 18, 64]. We first used automated annotation tools to assign functions to proteins. Because the annotations from these tools were not always accurate, we manually curated the annotations of 3,959 proteins and then extrapolated the functions to 14,547 similarly annotated proteins, which in total represented between 74 and 86 percent of the total microbial protein abundance in each sample (Extended Data Table 6). Based on the annotations, we assigned broad functional categories, such as amino acid metabolism, gene expression, glycan degradation, and monosaccharide metabolism and more detailed functional categories, such as ribosomal proteins and glycolysis to each of these proteins (Extended Data Table 6; Supplementary Fig. 3; Supplementary Results Section B). We then used the relative protein abundances to determine the investment of gut microbes into each of these functions. All of the broad functional categories, except for secondary metabolism, had significant changes in abundance due to dietary protein (ANOVA, *P* value < 0.05; Extended Data Table 7; Fig. 2; Supplementary Fig. 4), which indicates that the source of dietary protein changes the gut microbiota’s metabolism and physiology. Hierarchical clustering of all samples by abundances of broad functional categories revealed that the yeast and egg white diets clustered separately from all the other diets (Supplementary Fig. 5), similar to the results from the taxonomic clustering (Fig. 1g); however, a similar analysis at the detailed functional level revealed separate yeast, rice, and egg white clusters, with some outliers (Supplementary Fig. 6).

**Figure 2:**
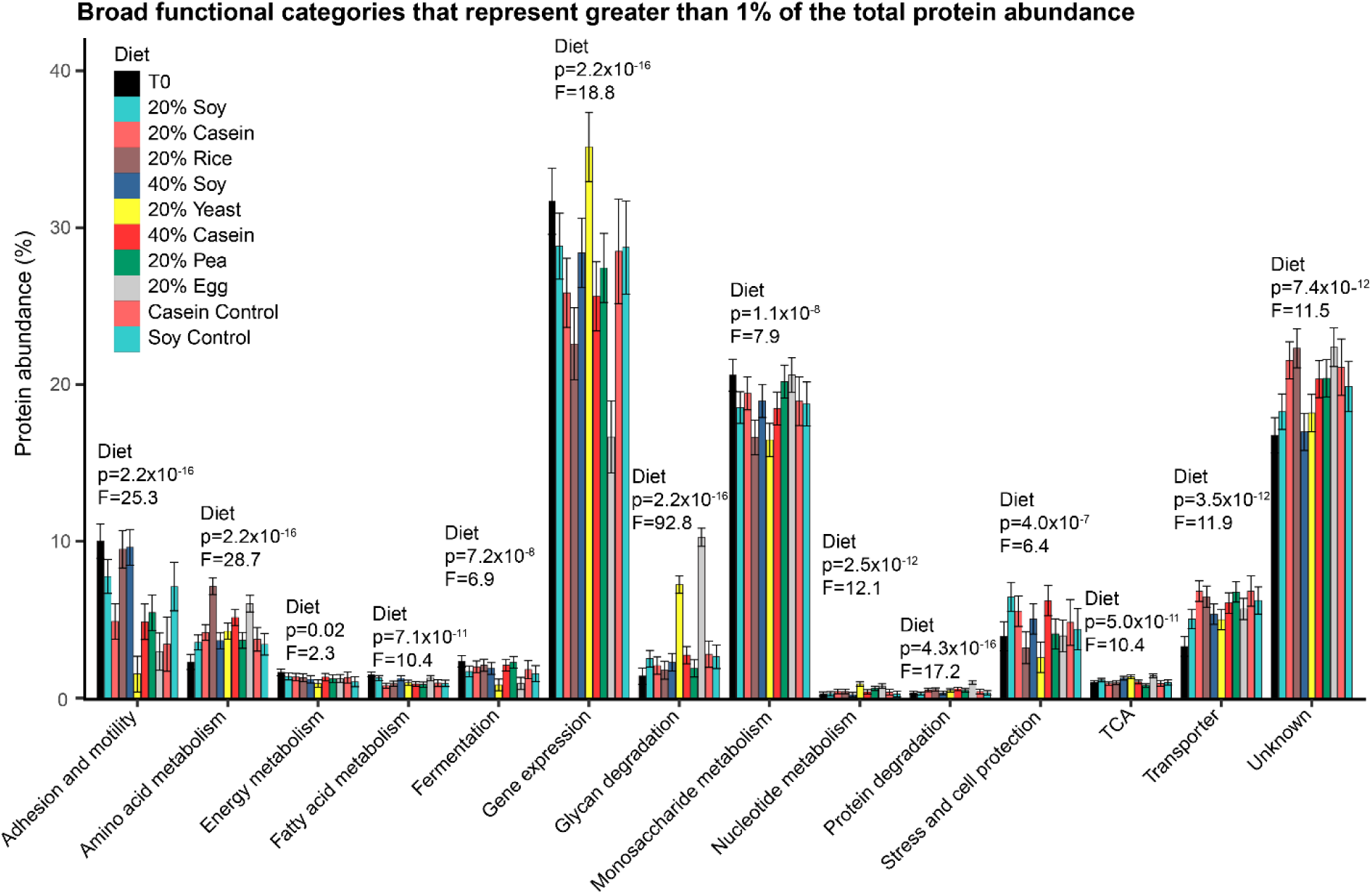
Broad functional categories of microbial proteins change significantly in abundance due to the source of dietary protein. Abundance of broad functional categories that represent at least 1% of the microbial protein abundance in at least one diet. The abundance is a modeled mean calculated from mixed effects models and the error bars represent 95% confidence intervals calculated from these models. All the categories shown here had a *P* value for the diet factor below 0.05; *P* value < 2.2x10^-16^ is the lower limit of the method. For underlying data see Extended Data Tables 7 and 8. For higher resolution of functional categories, e.g. fermentation, see Supplementary Figs. 3-4.

The two abundant broad functional categories (>1% of the total protein biomass) that had the greatest effect size due to diet were amino acid metabolism and glycan degradation, with *F* values of 29 and 93 respectively (Fig. 2; Extended Data Table 7). Amino acid metabolism increased in the brown rice and egg white diets relative to all other diets except the 40% casein diet and glycan degradation significantly increased in yeast and egg white diets relative to all other diets (Fig 2). Significant changes in the abundance of amino acid metabolism supported our initial expectation that the response of gut microbiota to different dietary protein sources would likely relate to amino acid metabolism; however, we also found that the abundance of glycan degrading enzymes responded more strongly to the source of dietary protein than did enzymes for amino acid metabolism. This suggests that glycan degradation instead of amino acid metabolism may be the major driver of taxonomic and functional changes in the gut microbiota in response to dietary protein source. We discuss these two functions in detail in subsequent sections. In addition, we observed specific changes in the abundance of enzymes associated with the gene expression, monosaccharide metabolism, fermentation, and stress and cell protection functional categories (Supplementary Results Section B; Supplementary Fig 3).

### Source of dietary protein alters the abundance of amino acid degrading enzymes

We manually classified 911 proteins (Extended Data Table 9) representing 68 enzyme functions (Extended Data Table 10) according to their involvement in the degradation (Fig. 3a), synthesis (Fig. 3b), interconversion (Fig. 3c), or reversible (Fig. 3d) reactions of specific amino acid pathways (Supplementary Figs. 7-18). In all diets except the yeast and standard chow diets, we observed that gut microbiota trended towards amino acid degradation instead of synthesis. We found that amino acid degrading enzymes were on average 2- to 6-fold more abundant than amino acid biosynthesis enzymes (Fig. 3a and 3b). Amino acid degrading enzymes were significantly more abundant in the rice and egg diets as compared to all the other diets (Fig. 3a), which is consistent with the observation that dietary proteins were significantly more abundant in the fecal samples of the brown rice and egg diets as compared to all other diets (Fig. 3e), suggesting that there may be a connection between the digestibility of dietary protein and amino acid degradation by the gut microbiota. Though amino acid synthesis enzymes were generally less abundant, we did observe a trend towards an increase in amino acid synthesis enzymes in the yeast protein diet relative to the other diets. This trend was not significant (ANOVA, *P* value = 0.09; Extended Data Table 11;Fig. 3b), but we observed several individual synthesis enzymes to be significantly increased in the yeast protein diet relative to other diets. These enzymes were involved in the synthesis of branched-chain amino acids (Suppl. Fig. 8), cysteine (Suppl. Fig. 9), lysine (Suppl. Fig. 15), proline (Suppl. Fig. 17), or tyrosine (Suppl. Fig 18).

**Figure 3:**
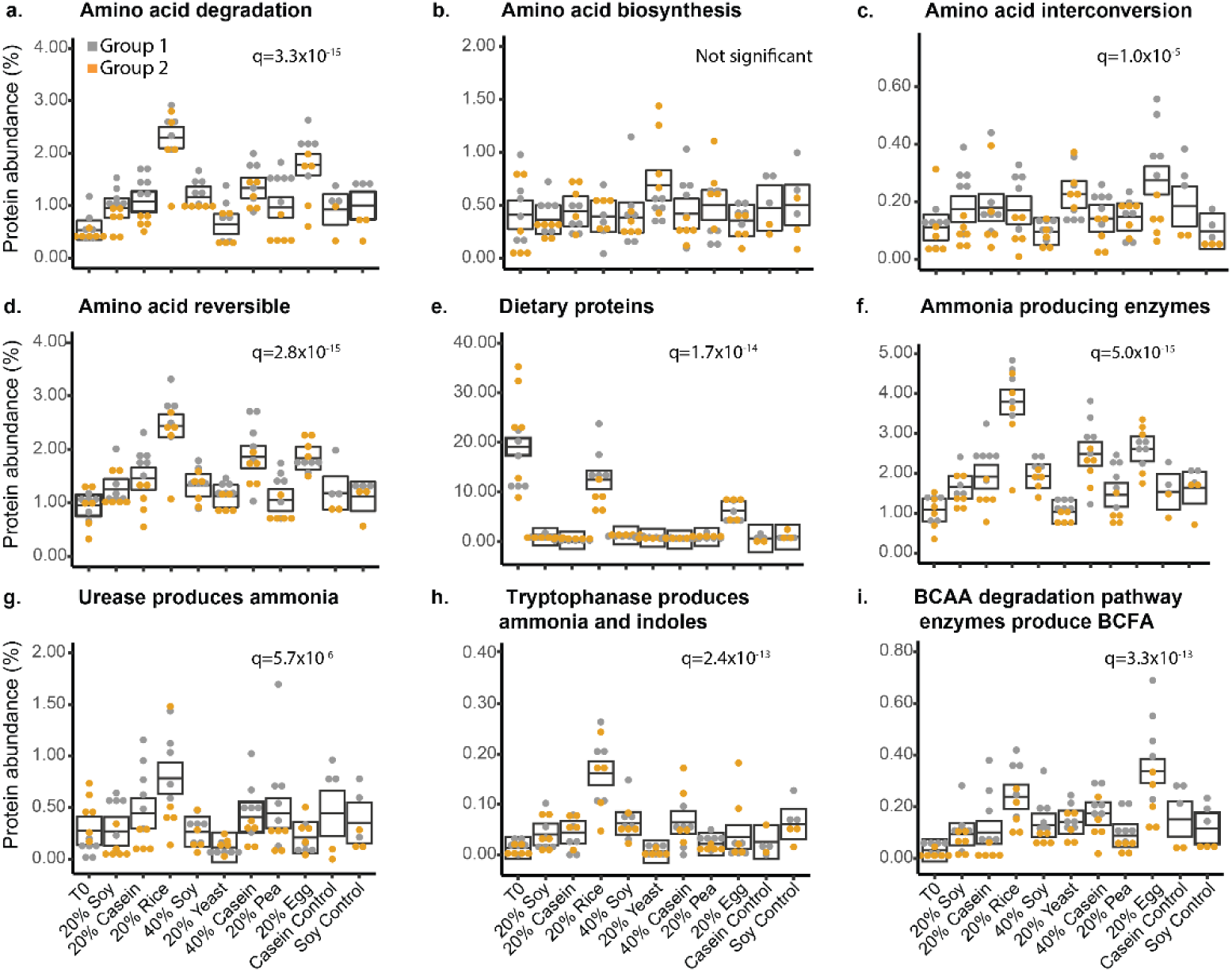
Amino acid degradation increases in the rice and egg diets. Box plots depict the percent abundance (out of the total microbial protein abundance) of different categories of microbial amino acid metabolism proteins. The exception is the dietary proteins, which are based on the percent protein abundance of the total metaproteome. The boxes represent the 95% confidence interval of the mean (line) for each diet from a complete mixed-effects model, and the q-values represent the FDR controlled *P* values for the diet factor from an ANOVA on these models (q<0.05 indicates significance)(Extended Data Tables 11-12). Boxes that do not overlap indicate statistical significance. Circle dots represent actual values per sample and are colored by mouse group. Abundance of proteins classified as (a) degrading an amino acid, (b) synthesizing an amino acid, (c) converting between two amino acids, and (d) reversible. (e) Abundance of all dietary proteins detected in each condition. (f) Abundance of enzymes that are likely to produce ammonia. Enzymes classified as degrading or reversible were included as long as ammonia was one of the potential products. Summed abundance of all proteins classified as (g) urease, (h) tryptophanase, and (i) involved in branched-chain amino acid (BCAA) degradation to branched-chain fatty acid (BCFA) (includes branched-chain amino acid aminotransferase or ketoisovalerate oxidoreductase).

Not all amino acid degrading enzymes increased in both the brown rice and egg white diets; sometimes they increased in one or the other (Supplementary Figs. 7-18). For example, enzymes associated with the degradation of threonine were more abundant in the egg white diet (Supplementary Fig. 12), whereas enzymes associated with tryptophan degradation were increased in the brown rice diet (Supplementary Fig. 18). Brown rice and egg were not the only diets in which the abundances of specific amino acid degrading enzymes increased. Alanine dehydrogenase increased in the 40% soy diet relative to the pea, yeast, 20% soy, and 20% casein diets (Supplementary Fig. 9) and cysteine desulfurase increased in the 40% casein and casein control diets relative to most other diets (Supplementary Fig. 9).

Changes in amino acid degradation by the gut microbiota have potential implications for host health by directly affecting local tissues or through interactions along the gut-brain axis depending on the metabolites produced by amino acid degradation pathways or the enzyme pathways themselves [6, 65]. For example, ammonia produced by amino acid deamination and hydrogen sulfide produced from methionine and cysteine degradation are toxic [66–68], indoles produced by tryptophanase and γ-aminobutyric acid (GABA) produced by glutamate decarboxylase are neurotransmitters [69, 70], branched-chain fatty acids produced by branched-chain amino acid degradation have been suggested to be anti-inflammatory [71, 72], and proline metabolism has been linked to the depression [73] and enteric infections [74]. We found ammonia producing enzymes to be significantly more abundant in the brown rice diet as compared to all other diets, and also more abundant in the egg white and 40% casein diets as compared to the standard chow, 20% soy, yeast, pea, and control diets (Fig. 3f and 3g).

Tryptophanase significantly increased in the brown rice diet relative to all other diets, whereas glutamate decarboxylase increased in the egg white diet relative to all other diets except brown rice, pea, and the control diets (Fig. 3h and Supplementary Fig. 7). We observed that branched-chain amino acid degrading enzymes were significantly increased in the egg white protein diet relative to all other diets (Fig. 3i), and proline degrading enzymes were increased in the brown rice diet relative to other diets, except the 40% soy and 40% casein diets where we also observed proline degradation to be significantly increased relative to the standard chow, yeast, and pea diets (Supplementary Fig. 17). These results show that the source of dietary protein can alter overall amino acid metabolism in the gut microbiome, as well as the abundance of different pathways. These changes have the potential to affect host physiology and health.

### Gut microbes express distinct glycoside hydrolases to grow on different sources of dietary protein

Glycan degrading enzymes (glycoside hydrolases) showed the largest overall changes in response to dietary protein source (Fig. 2, Extended Data Table 7). Specifically, these enzymes increased significantly in abundance in the yeast and egg white diets compared to the other diets. To further investigate the interaction of these glycan degrading enzymes with dietary protein we manually curated the functional assignments and potential substrate specificity of the 1,059 microbial glycoside hydrolases detected in our metaproteomes (Extended Data Table 13).

We grouped the validated glycoside hydrolases into 91 families based on the CAZy database (Extended Data Table 14) [53]. Of these families, 54 significantly changed in abundance between the different dietary protein sources (ANOVA, q<0.05) (Extended Data Table 15). Different glycoside hydrolase families increased in abundance in the soy, casein, brown rice, yeast, and egg white diets suggesting that distinct glycans drive their abundance changes across the different diets (Fig. 4a, Extended Data Table 14-16, Supplementary Results Section C). The most abundant glycoside hydrolase families, GH18 in the case of egg white and GH92 in the case of yeast, have previously been associated with the degradation of glycans conjugated onto proteins (glycosylations) as part of polysaccharide utilization loci (PULs). PULs are operons that contain all the proteins necessary to import and degrade a specific glycan structure [75]. These GH18s are endo-β-N-acetylglucosaminidases that break the bond between two acetylglucosamine residues attached to asparagine in N-linked glycoproteins. This reaction releases the glycan from the glycoprotein [76]. Meanwhile, GH92s, which are alpha-mannosidases, have been previously associated with the release of mannose residues from the glycosylations on yeast mannoproteins [76].

**Figure 4:**
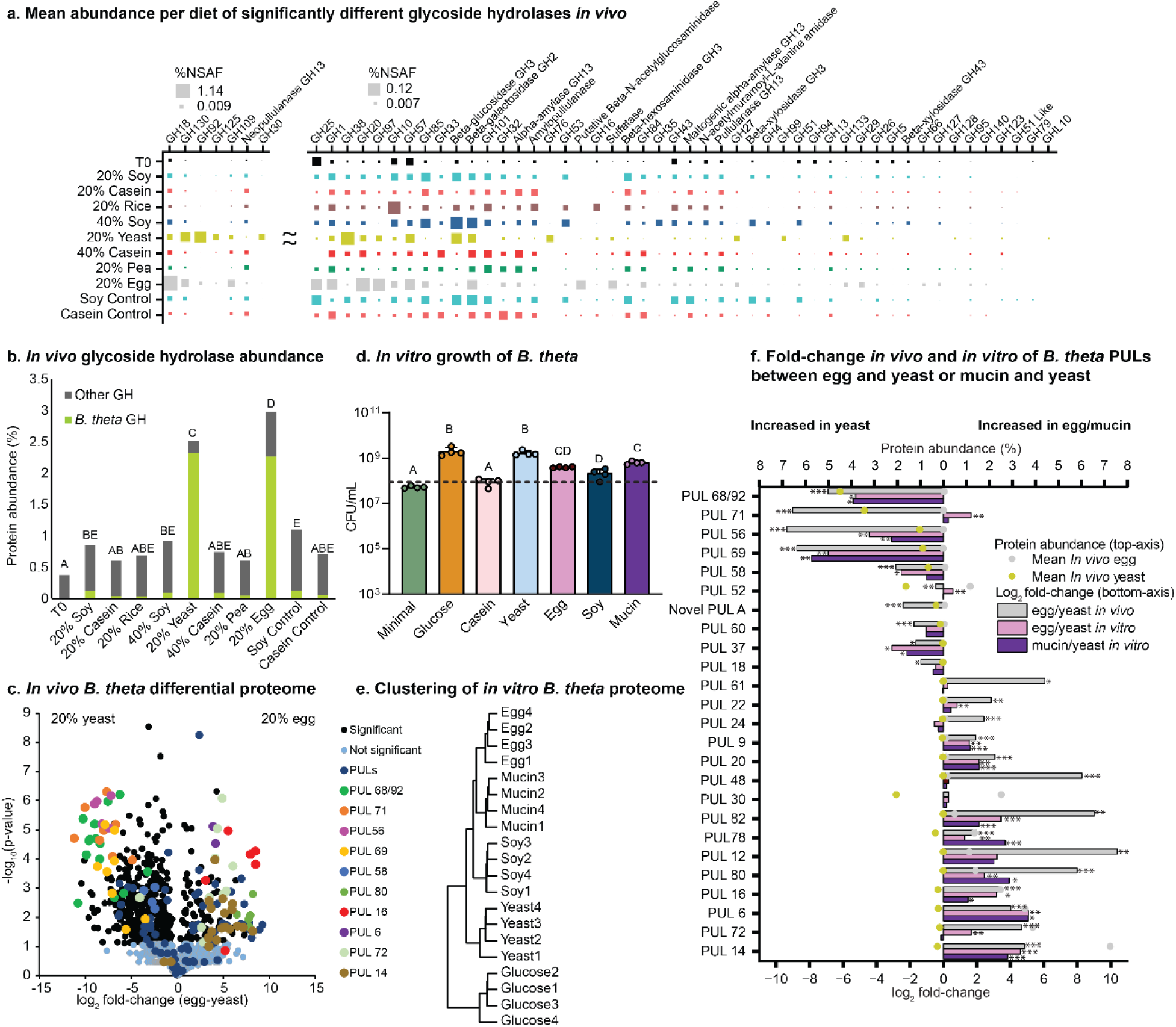
Glycosylations on dietary proteins drive shifts in microbial composition. (a) Mean summed protein abundance per diet of glycoside hydrolases with significantly different abundances between diets (q<0.05 in mixed-effects ANOVA models). (b) Mean combined protein abundance of proteins predicted to be glycoside hydrolases. The proportion of these proteins that belong to *B. thetaiotaomicron* is highlighted. Diets that do not have overlapping letters also have non-overlapping 95% confidence intervals for each diet calculated from a complete mixed-effects model. (c) Volcano plot of -log10 *P* values (Welch’s t-test; FDR controlled at q<0.05) versus the log2 fold-change of *B. thetaiotaomicron* proteins under the yeast and egg white protein diets *in vivo* after recalculating the protein abundance based on proteins only assigned to *B. thetaiotaomicron*. Filled circle symbols, indicating individual proteins, were colored based on the polysaccharide utilizing locus (PUL) operon to which the protein belongs. We only colored the proteins from PULs that had an absolute difference of 0.5% or greater between the yeast and egg diets. (d) Colony forming units per mL (CFU/mL) of *B. thetaiotaomicron* grown in defined media with dietary proteins as the sole carbon source. The dotted line indicates T0 CFU/mL. Media that do not share letters are significantly different based on ANOVA and Tukey HSD multiple comparisons after log transformation (*P* value < 0.05). (e) Hierarchical clustering (ward.D2 on Euclidean distances) of the *in vitro B. thetaiotaomicron* proteome under different media. (f) *In vivo* and *in vitro* comparison of the summed protein abundance of PULs. The bottom axis depicts the log2 fold-change between egg white and yeast protein or mucin and yeast protein. The top axis depicts the mean protein abundance of the PULs *in vivo* in the yeast diet on the left and in the egg diet on the right. A Welch’s t-test (with FDR control) was performed between each comparison to detect significant changes in PUL protein abundances (*** = q < 0.01, ** = q < 0.05, * = q < 0.1) (Extended Data Tables 18 and 20).

We found that the total abundance of glycoside hydrolases increased from <1% in the majority of diets to >2.5% in the yeast and egg diets (Fig. 4b). Additionally, we observed a general trend towards an increased abundance of glycoside hydrolases in all defined diets compared to the T0 (standard chow) diet; however, the increase was only significant for the soy, yeast and egg diets (Fig. 4b). The majority of the glycoside hydrolases in the yeast and egg diets came from *B. thetaiotaomicron* (Fig. 4b). Because *B.thetaiotaomicron* is one of the primary drivers of the changes in microbiota composition in these diets (Fig. 1g), this suggests that glycoside hydrolases are closely associated with the observed changes in microbiota composition.

To examine the specific role of *B. thetaiotaomicron*in glycan degradation in the yeast and egg white diets, we compared the abundances of all *B. thetaiotaomicron* proteins in the metaproteome between the two diets. Out of 1,420 detected *B. thetaiotaomicron* proteins, the abundances of 592 proteins significantly differed between the two diets (q < 0.05, Welch’s t-test) (Fig. 4c; Extended Data Table 17). Many of the significant proteins that were the most abundant and had the greatest fold-change between the two diets came from PULs (Fig. 4c; Supplementary Fig. 19). Between 10% and 25% of the total protein abundance of *B. thetaiotaomicron*in in the yeast and egg white diets came from these PULs (Extended Data Table 18). The proteins belonging to each PUL tended to be expressed together either being significantly increased in egg white or the yeast diet (Figure 4c).

Several of the PULs that increased when we fed mice the yeast diet have previously been shown to specifically degrade the glycosylations on yeast cell wall proteins. PULs 68/92 (BT3773-3792) and 69 (BT3854-3862) (Fig. 4c; Supplementary Fig. 19) degrade ⲁ-mannans attached to yeast mannoproteins in *Saccharomyces cerevisiae* [76], whereas PUL 56 (BT3310-3314), degrades yeast β-glucans also attached to yeast cell wall mannoproteins [77]. Conversely, the majority of the PULs increased when we fed mice the egg white diet had been previously linked to growth on mucin glycan conjugates: PUL14 (BT1032-1051), PUL6 (BT3017-0318), PUL16 (1280-1285), PUL 80 (BT4295-BT4299), and PUL12 ( BT0865-0867) (Supplementary Fig. 19) [78]. An additional abundant PUL, PUL72 (BT3983-BT3994), has been previously implicated in the degradation of mannoproteins of mammalian origin [76] and our result suggests that PUL72 is also involved in the degradation of mannoproteins from non-mammalian vertebrates.

To test if *B. thetaiotaomicron* could grow on yeast and egg white protein as predicted from the *in vivo* data, and if the expression of PULs was driven by direct responses to the dietary protein sources, we characterized *B. thetaiotaomicron* growth and its proteome on dietary protein sources *in vitro*. We used a defined culture media and supplemented purified dietary protein sources as the sole carbon source to determine if this supported *B. thetaiotaomicron* growth. We found that *B. thetaiotaomicron* grew in the presence of glucose (control), yeast protein, egg white protein, soy protein, and intestinal mucin (Fig. 4d, Tukey HSD adj P < 0.05). We analyzed the proteomes of *B. thetaiotaomicron* in these five different conditions *in vitro* to determine if PULs played a role in growth (Extended Data Table 19). An overall comparison of the proteome between the media supplemented with 4 different protein sources revealed that egg white protein and mucin had the most similar proteome, and the proteome from the glucose control clustered separately from the protein-sources (Fig. 4e; Supplementary Fig. 20). We observed that 15 out of 24 PULs that were significantly different in abundance between the egg white and yeast diets *in vivo* were also significantly different in the same direction *in vitro* (Fig. 4f). In addition, 12 of these 15 PULs showed the same expression pattern in both the mucin and egg white protein media as compared to the yeast protein medium (Fig. 4f, Extended Data Tables 18 and 20). The relationship between mucin and egg white metabolism in microbiota species *in vivo* is further supported by the fact that five of the six species with greater than 5% abundance in an egg white sample (*B. thetaiotaomicron*, *A. muciniphila*, *Atopobiaceae* bacterium AB25_9 *Paramuribaculum sp.*, *D. newyorkensis*) had abundant enzymes associated with the metabolism of sugars usually thought to be derived from mucin. These enzymes, of which several were among the top 100 most abundant proteins of these organisms, catalyze the metabolism of sialic acid (N-acetylneuraminate lyase, N-acylglucosamine 2-epimerase), N-acetylglucosamine (N-acetylglucosamine-6-phosphate deacetylase, glucosamine-6-phosphate deaminase, PTS system N-acetylglucosamine-specific, or fucose (fucosidase, fucose isomerase) (Extended Data Table 6). In summary, these results indicate that the glycosylations on yeast and egg white proteins drive the increase in abundance of *B. thetaiotaomicron* in the yeast and egg white diets, and that egg white proteins and intestinal mucin share similar glycosylations leading to the expression of similar PULs for their degradation.

## DISCUSSION

In this study, we sought to characterize how dietary protein source affects the gut microbiota’s composition and function by measuring species-resolved proteins using integrated metagenomics-metaproteomics. We showed that source of dietary protein significantly alters the gut microbiota’s composition, more so than amount of protein, and that yeast and egg white protein had the greatest effect on the composition driven by an increase in the relative abundance of *B. thetaiotaomicron* and a decrease of bacteria from the class *Clostridia*. We also showed that the source of dietary protein altered the overall functional profile of the gut microbiota as reflected by changes in the abundance of microbial proteins assigned to broad functional categories. In particular, proteins involved in amino acid metabolism increased in abundance in the brown rice and egg white diets, whereas enzymes assigned to glycan degradation increased in the yeast and egg white diets.

The increase in amino acid metabolizing enzymes in the brown rice and egg white diets was driven by amino acid degrading enzymes. Previous studies across multiple species have shown that increasing the amount of protein fed to animals leads to an increase in the ammonia concentration in stool [79–81], which suggests that increased protein availability leads to increased amino acid deamination or urease activity in the gut. Here we show that, regardless of the amount of protein, the source of protein itself can lead to increases in amino acid deaminating enzymes and ureases from the intestinal microbiota. Gut microbiota urease activity and amino acid deamination have been linked to serious diseases like hepatic encephalopathy when liver function is disrupted [82]. Replacement of bacteria that produce these deaminating enzymes and ureases with bacteria that do not has been suggested as a potential treatment [83], our results suggest that adjustments in dietary protein source could be considered as well.

Because amino acids are the backbone of protein, we expected to observe changes in the abundance of amino acid degrading enzymes between the different sources of dietary protein; however, the effect of dietary protein source on the abundance of glycan degrading proteins was even greater than the effect on amino acid degrading enzymes. Our results suggest that the increase in glycan degrading proteins in the yeast and egg white diets is due to the glycosylations conjugated to these proteins. Yeast and egg white proteins have distinct glycan conjugate structures [84–86]. In the presence of yeast dietary protein we were able to show, *in vivo* and *in vitro,* increased expression of PULs associated with the degradation of yeast mannoprotein glycan conjugates. In the presence of egg white protein, we observed an increase in PULs previously linked to the degradation of the glycan conjugates of mucin. This combined with increases in mucin foraging bacteria *Akkermansia muciniphila* [61] and *Paramuribaculum* sp. [62] suggests that egg white protein promotes the abundance of mucin foraging bacteria and their proteins. The link between the foraging of mucin and egg white protein in retrospect makes sense, as egg white protein contains mucins called ovomucin and other proteins: ovalbumin, ovotransferrin, and ovomucoid, which have been previously shown to be N-glycosylated with acetylglucosamine and mannose containing glycans [84, 87]. Previous studies in mice have shown that diets, which promote bacteria and their enzymes that degrade mucins, can make the host more susceptible to enteric inflammation and infection [72, 88]. Because egg white protein also promotes these functions, these results suggest that diets high in egg protein may be detrimental to gastrointestinal health, which could explain the prior results from population level studies that eggs lead to increased mortality rates among humans [3].

Our study has at least two limitations preventing direct translation of microbiota responses to dietary protein sources into a human health context. First, we used purified dietary proteins, which differ from commonly consumed dietary proteins in that regular dietary protein sources also provide some amount of additional major dietary components such as fats, carbohydrates, and fiber. For example, plant proteins usually come with a relevant amount of fiber, whereas animal proteins are often low in fiber and have higher content fats [8]. Second, we used fully defined diets which allowed us to track effects to specific protein sources, but we anticipate that the dietary context of protein sources such as co-consumption of multiple protein, fiber, fat and carbohydrate sources will strongly influence the interactions of dietary protein sources with gut microbiota. For example, these diets only had cellulose as the fiber source, which cannot be metabolized by many microbes that are commonly considered fiber degraders. Thus, the low diversity in fiber sources of these diets likely amplified the effects of yeast and egg white protein in this study. The power of our study lies in our ability to confirm that the source of dietary protein does impact gut microbiota function and should be considered when thinking about how diet impacts gut microbiota and its implications for host health. Future studies that determine how the effect of dietary protein source on the gut microbiota impacts gastrointestinal diseases are needed.

## Declarations

### Ethics approval and consent to participate

We conducted our animal experiments in the Laboratory Animal Facilities at the NCSU CVM campus (Association for the Assessment and Accreditation of Laboratory Animal Care accredited), which are managed by the NCSU Laboratory Animal Resources. Animals assessed as moribund were humanely euthanized via CO2 asphyxiation, as approved by NC State’s Institutional Animal Care and Use Committee (Protocol # 18-034-B).

## Data availability

The mass spectrometry proteomics data have been deposited to the ProteomeXchange Consortium via the PRIDE [89] partner repository with the dataset identifier PXD041586 (metaproteomic data) and PXD050296 (*B. thetaiotaomicron* in vitro proteomics data). Metagenomic raw reads were submitted to NCBI SRA under the bioproject identifier PRJNA1026909. All metagenome assembled genomes (MAGs) with accompanying metadata were submitted to DRYAD DOI: 10.5061/dryad.x0k6djhq5. [Reviewer link: https://datadryad.org/stash/share/QagcDe_b_b0GbbyQ7mPOxBapFL3QbaXt3-fhiZRvDCM]

## Acknowledgments

We thank Brandon C. Iker for inspiring this research and initial concepts, Tjorven Hinzke for advice on statistics, Heather Maughan for editing of the manuscript, Erin Baker for advice on methods, and Lawrence David and Balfour Sartor for advice and consultation on the project. All LC-MS/MS measurements were made in the Molecular Education, Technology, and Research Innovation Center (METRIC) at North Carolina State University.

## Author Contributions

JABR: Experimental design, data collection, data processing, data analysis, author of original manuscript, editing

AB: Conceptualization of the study, experimental design, data collection, data processing, editing

ASM: Experimental design, data collection, editing AA: Data processing, analysis, editing

MVW: Data processing, analysis, editing

AKM: Data processing

JM: Coding, graphic design

SV: Data collection

TR: Data collection

CMT: Experimental design, data collection, editing

MK: Funding, conceptualization of the study, experimental design, data collection, data processing, data analysis, writing, editing

