## Supplemental Figures for "Dietary protein source alters gut microbiota composition and function"

### Supplementary Results

#### Section A: Effect of dietary protein on gut microbiota composition

Hierarchical clustering of the proteinaceous biomass of species revealed distinct starting microbiota compositions for the group 1 and group 2 mice. This led us to surmise that the source of dietary protein altered the gut microbiota regardless of starting microbiota composition. To confirm this, we analyzed the group 1 and group 2 mice separately. We observed that they had similar alpha diversity and richness responses to dietary protein source regardless of mouse group and that the Bray-Curtis dissimilarity patterns were also similar (Fig. 1d and e; Supplementary Fig. 1a, b and c). Principal component analysis showed the same separation as the hierarchical clustering between mouse group 1 and 2 along the first component (Supplementary Fig. 1d). Further analysis along the second and third components showed distinct dietary clusters between 1) soy, 2) T0, 3) yeast and egg white, and 4) casein, brown rice and pea (Supplementary Fig. 1e). Principal component analysis on the separate mouse groups revealed the same cluster groups (Supplementary Fig. 1f and g), suggesting that the source of dietary protein alters the gut microbiota's composition regardless of starting microbiota composition.

We observed dynamic species abundance responses to different sources of dietary protein; these responses were in some cases consistent across mouse groups, while in other cases they were mouse group specific. For example, *Bacteroides thetaiotaomicron* (*B. theta*) increased in abundance across both mouse groups in the yeast and egg white diets (Fig. 1g; Supplementary Fig. 2), while the abundances of *Schaederella* sp. AB67-1 and Lachnospiraceae bacterium AB103-0 repeatedly increased in the presence of soy (Supplementary Fig. 2). Other changes in species abundance due to diet were mouse group specific. For example, *Lactobacillus johnsonii* increased in abundance in the pea and casein diets in the group 1 mice, while *Faecalibaculum rodentium* showed a similar pattern in the group 2 mice. *Oscillospiraceae* bacterium AB63-2 increased in abundance in the presence of the soy diets in the group 1 mice, while *Oscillospiraceae* bacterium AB54-6 followed a similar pattern in the group 2 mice. All of the abundant species were significantly different in abundance between at least two diets (Supplementary Fig. 2). Supplementary Fig. 2 contains the details for the specific dynamics of the 36 most abundant species we detected. All species had a significantly different abundance between diets (linear mixed effect model ANOVA, $q < 0.05$).

#### Section B: Effect of dietary protein on microbiota function

The two most abundant broad functional categories of detected peptides were gene expression, which includes ribosomes, chaperones, and transcription related enzymes, and monosaccharide metabolism, which includes glycolysis and the metabolism of simple sugars other than glucose (Fig. 2). The microbial investment in gene expression enzymes increased in the yeast diet relative to all other diets (except standard chow) and decreased in the egg white diet relative to all other diets. This was driven by an overall increase in the abundance of ribosomal proteins in the yeast diet (Supplemental Fig. 3b). The abundance of ribosomal proteins has been suggested to be directly correlated with bacterial growth rates[1] suggesting that overall bacterial growth rate is higher when mice were fed the yeast diet. This is further supported by the overall higher bacterial load in the yeast diet (Fig. 1c). In contrast, we observed gene expression proteins that assist with the synthesis and folding of proteins, e.g., elongation factors and chaperones, to be increased in the soy diets relative to some of the other diets (Supplemental Fig. 3b). Curiously, we also observed an increase in stress proteins, including oxidative stress proteins, in the soy and casein diets relative to brown rice, egg white, and yeast (Supplementary Fig. 3c). Oxidative stress interferes with the proper elongation and folding of proteins, which could explain why chaperones and elongation factors are increased in the soy diet[2].

We observed a significant decrease in the abundance of monosaccharide metabolizing enzymes in the yeast and brown rice diets (Fig. 2). Most abundant in this category were the enzymes belonging to the energy pay-off phase of glycolysis (Supplementary Fig. 3d). However, many of the other functions within monosaccharide metabolism had different abundance patterns. For example, galactose and mannose metabolism enzymes were increased in the yeast diet (Supplementary Fig 3e), while along with galactose metabolism, we also observed a general increase in the abundance of fucose, glucosamine, and sialic acid metabolism in the egg white diet relative to other diets. Glucosamine and sialic acid metabolism were also increased in the casein and pea diets relative to other diets (Supplementary Fig 3e). Fucose, galactose, sialic acid, and acetylglucosamine are all components of mucin[3]. In summary, these results suggest that the source of dietary protein impacts sugar metabolism in the gut microbiota.

Two other broad functional categories that significantly changed in abundance due to dietary protein source were adhesion and motility proteins and fermentation proteins. The microbiota invested significantly less in proteins categorized as adhesion and motility proteins in the yeast and egg diets. Flagellar proteins drove this result, which can be explained by the replacement of species from the class Clostridia with species from the class Bacteroidia because microbes in the phylum Bacteroidota usually do not have flagella[4]. The microbial investment in fermentation enzymes also decreased in the yeast and egg white diets. We divided fermentation enzymes into three categories, ethanol producing, short-chain fatty acid (SCFA) producing, and lactic acid producing (Supplementary Fig. 3a). This categorization revealed that fermentation enzymes leading to SCFA metabolites were the primary drivers of the decrease in fermentation-related proteins in the yeast and egg white diets. Production of SCFAs has been previously linked to anti-inflammatory responses, which could suggest that the changes in microbiota composition observed in the yeast and egg white diets may be detrimental to host health[5,6].

**Section C: Effect of brown rice and soy dietary protein on glycoside hydrolase abundance**

Several glycoside hydrolases increased in the presence of the brown rice and soy diets. In the soy diet, the expression of glycoside hydrolases was reproducible, increasing in abundance each of the three times the mice were fed a soy diet. Most notably β-glucosidases and β-xylosidases from the CAZy protein family GH3 were increased in the soy diets, while the abundance of GH16s increased in the brown rice diet. The abundance change of GH16s in the brown rice diet was driven by a single protein from the uncultured species *Oscillospiraceae* bacterium AB63-2 that was only detected in the brown rice diet. In the presence of soy protein, but not brown rice protein, this bacterium reproducibly expressed different glycoside hydrolases including two β-glucosidases and one β-xylosidase from the protein family GH3 (Extended Data Table 13). These results suggest that different protein sources generally affect the function and composition of the gut microbiota through the different glycan structures attached to their glycoproteins.

Supplementary Table 1: Composition of nine fully defined diets

| Diet | 20% Soy | 20% Casein | 20% Brown Rice | 40% Soy | 20% Yeast | 40% Casein | 20% Pea | 20% Egg White Solids, spray-dried | 20% Chicken bone broth |
|---|---|---|---|---|---|---|---|---|---|
| Teklad Catalog Number | TD.190249 | TD.190254 | TD.190252 | TD.190250 | TD.190253 | TD.190255 | TD.190251 | TD.190256 | TD.19025 7 |
| Protein supplier | Teklad | Teklad | Swanson | Teklad | Teklad | Teklad | NAKED | Teklad | Ancient Nutrition |
| Protein (g/Kg) | 230 | 230 | 260 | 460 | 400 | 460 | 222.22 | 248.45 | 220 |
| Sucrose (g/Kg) | 436.1 | 434.7 | 400.43 | 211.785 | 279.06 | 208.96 | 438 | 412.667 | 450.33 |
| Corn Starch (g/Kg) | 200 | 200 | 200 | 200 | 200 | 200 | 200 | 200 | 200 |
| Corn Oil (g/Kg) | 52.3 | 52.3 | 54.6 | 50 | 42.6 | 50 | 50.82 | 54.6 | 44.7 |
| Cellulose (g/Kg) | 37.86 | 37.86 | 37.86 | 37.86 | 37.86 | 37.86 | 37.86 | 37.86 | 37.86 |
| Vitamin Mix, Teklad (40060) (g/Kg) | 10 | 10 | 10 | 10 | 10 | 10 | 10 | 10 | 10 |
| Ethoxyquin, antioxidant | 0.01 | 0.01 | 0.01 | 0.01 | 0.01 | 0.01 | 0.01 | 0.01 | 0.01 |
| Mineral Mix, Ca-P Deficient 79055 (g/Kg) | 13.37 | 13.37 | 13.37 | 13.37 | 13.27 | 13.37 | 13.37 | 13.37 | 13.37 |
| Calcium Phosphate dibasic (g/Kg) | 15.26 | 16.66 | 23.72 | 6.8 | 0 | 9.6 | 19.72 | 22.54 | 23.73 |
| Calcium Carbonate (g/Kg) | 5.1 | 5.1 | 0 | 10.175 | 17.1 | 10.2 | 8 | 0.5 | 0 |

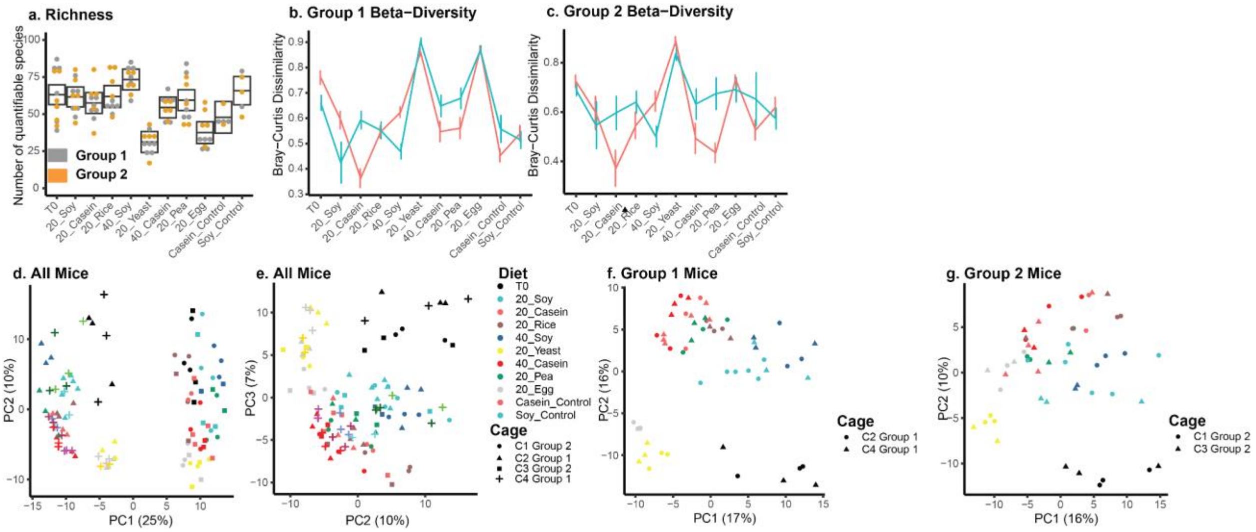


**Supplementary Figure 1: Source of dietary protein alters the gut microbiota's composition.** (a) depicts the average number of quantifiable species per diet; boxes represent the 95% confidence interval based on linear mixed effects models (Extended Data Table 2). (b) and (c) depict the Bray-Curtis dissimilarity between 20% soy diets (teal) or 20% casein diets (red) and all other diets for group 1 and group 2, respectively. (d) and (e) first, second, and third principal components of microbiota composition based on species level metaproteomic proteinaceous biomass. (f) and (g) first and second components of microbiota composition based on species level metaproteomics proteinaceous biomass for the group 1 and group 2 mice, respectively.

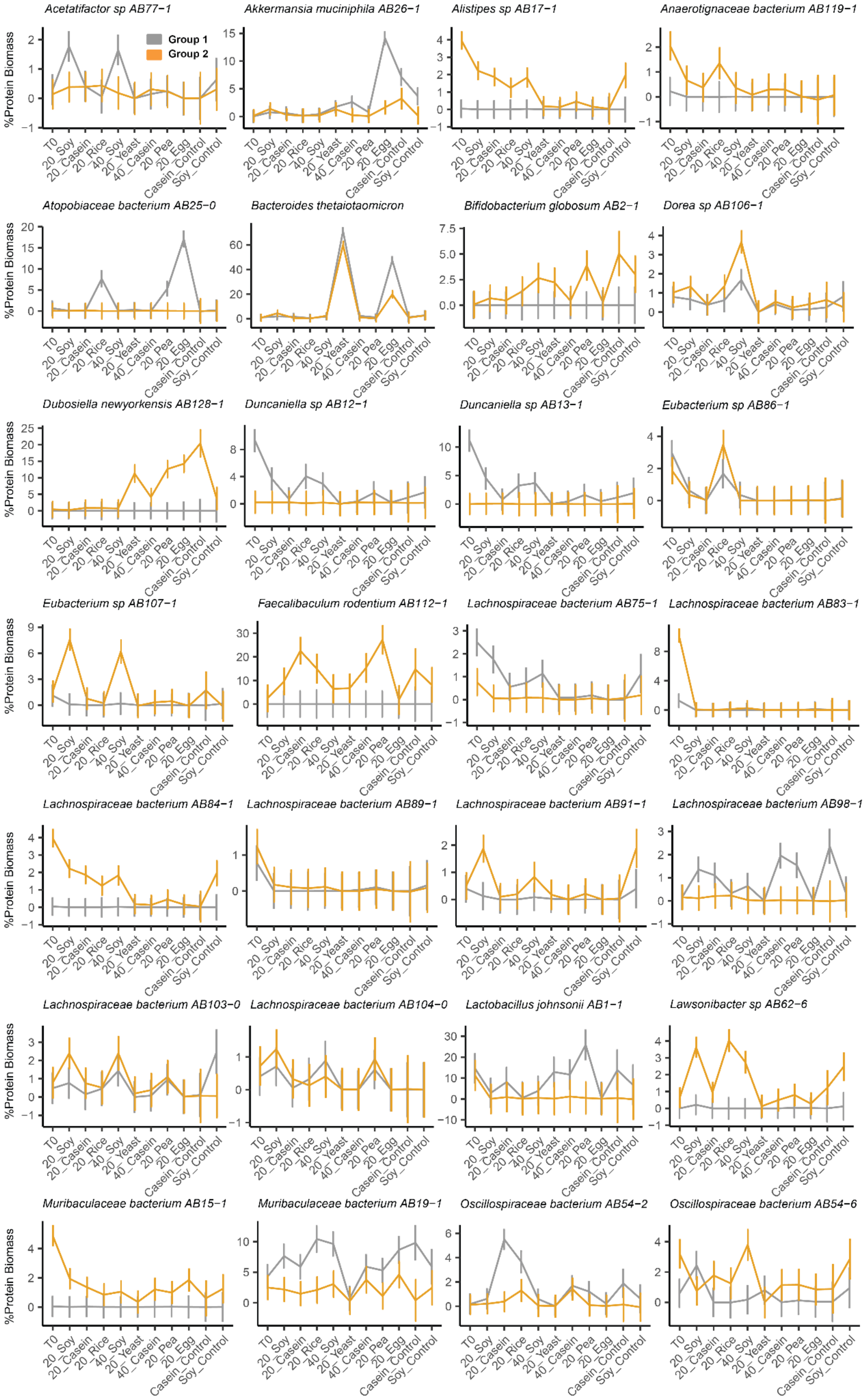
Acetatifactor sp AB77−1
Akkermansia muciniphila AB26−1
Alistipes sp AB17−1
Anaerotignaceae bacterium AB119−1
Atopobiaceae bacterium AB25−0
Bacteroides thetaiotaomicron
Bifidobacterium globosum AB2−1
Dorea sp AB106−1
Dubosiella newyorkensis AB128−1
Duncaniella sp AB12−1
Duncaniella sp AB13−1
Eubacterium sp AB86−1
Eubacterium sp AB107−1
Faecalibaculum rodentium AB112−1
Lachnospiraceae bacterium AB75−1
Lachnospiraceae bacterium AB83−1
Lachnospiraceae bacterium AB84−1
Lachnospiraceae bacterium AB89−1
Lachnospiraceae bacterium AB91−1
Lachnospiraceae bacterium AB98−1
Lachnospiraceae bacterium AB103−0
Lachnospiraceae bacterium AB104−0
Lactobacillus johnsonii AB1−1
Lawsonibacter sp AB62−6
Muribaculaceae bacterium AB15−1
Muribaculaceae bacterium AB19−1
Oscillospiraceae bacterium AB54−2
Oscillospiraceae bacterium AB54−6
Group 1
Group 2
%Protein Biomass
T0
20_Soy
20_Casein
20_Rice
40_Soy
20_Yeast
40_Casein
20_Pea
20_Egg
Casein_Control
Soy_Control

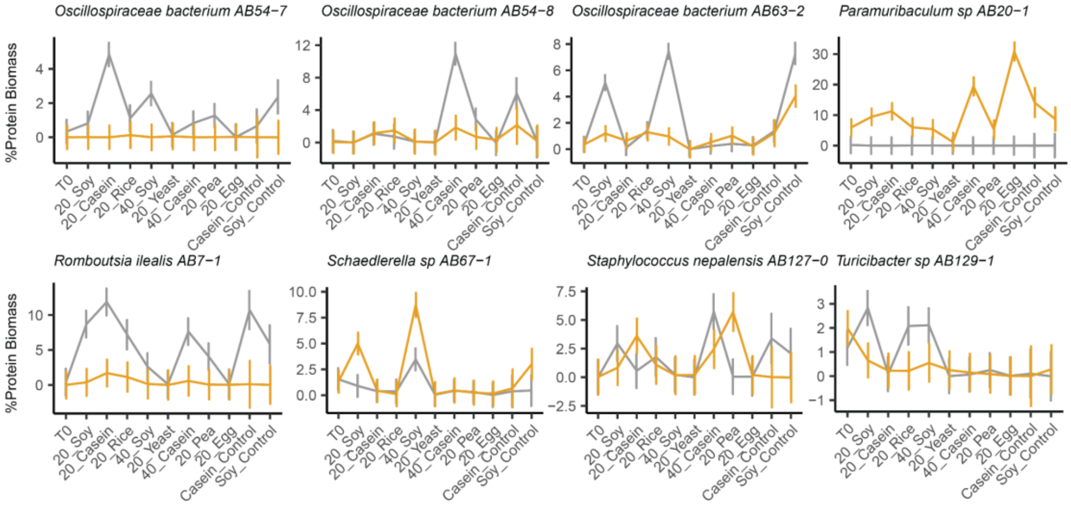


**Supplementary Figure 2: Abundances of the most abundant species across diets and mouse groups**. Line plots depicting the average abundance of bacterial species in the group 1 mice (gray) and group 2 mice (orange) across all diets. Abundances were determined from metaproteomic data using a biomass assessment method[7]. Error bars represent the 95% confidence interval of the mean using mixed effects modeling. Species were defined as abundant if they represented at least 5% of the microbial protein mass in one sample.

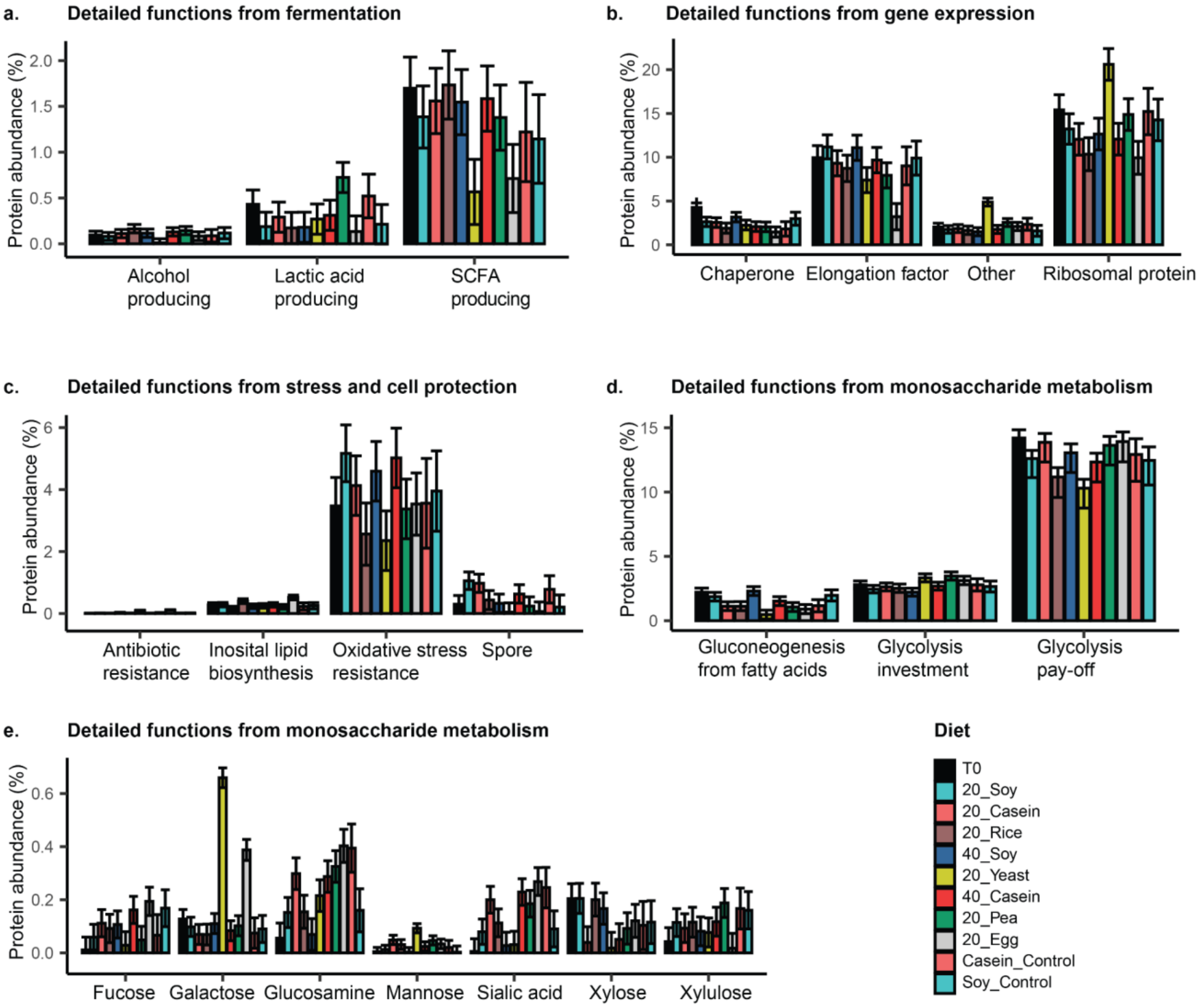


**Supplementary Figure 3: Abundance of detailed functional categories associated with fermentation, gene expression, stress and cell protection, and monosaccharide metabolism.** The mean protein abundance (% of total microbial proteins) per sample of each detailed function based on a complete linear mixed effects model. Error bars represent 95% confidence intervals and error bars that do not overlap indicate significant abundance differences. (a) Detailed functions that make up the fermentation broad functional category. (b) Detailed functions that make up the gene expression functional category. (c) Detailed functions that make up the stress and cell protection functional category. (d) The most abundant detailed functions that make up the monosaccharide metabolism functional category. (e) Select detailed functions that make up part of the monosaccharide metabolism category (Extended Data Tables 6, 7 and 8).

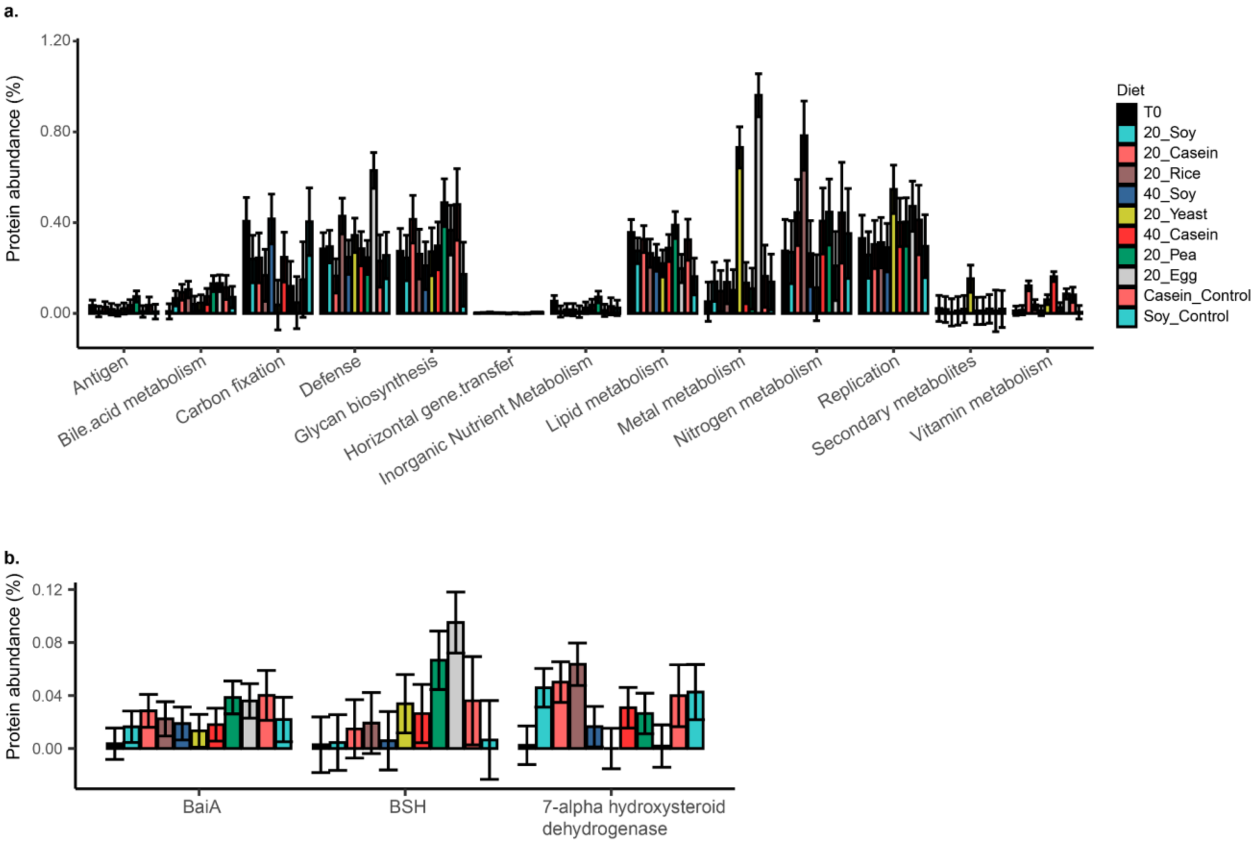


**Supplementary Figure 4: Effect of dietary protein source on lower abundance protein functions.** The mean protein abundance (% of total microbial proteins) per sample of each detailed function based on a complete linear mixed effects model. Error bars represent 95% confidence intervals and error bars that do not overlap indicate significant abundance differences. (a) Broad functional categories that represented less than 1% of the microbial protein abundance. (b) Consensus names of specific bile acid modifying enzymes (Extended Data Tables 6, 7 and 8).

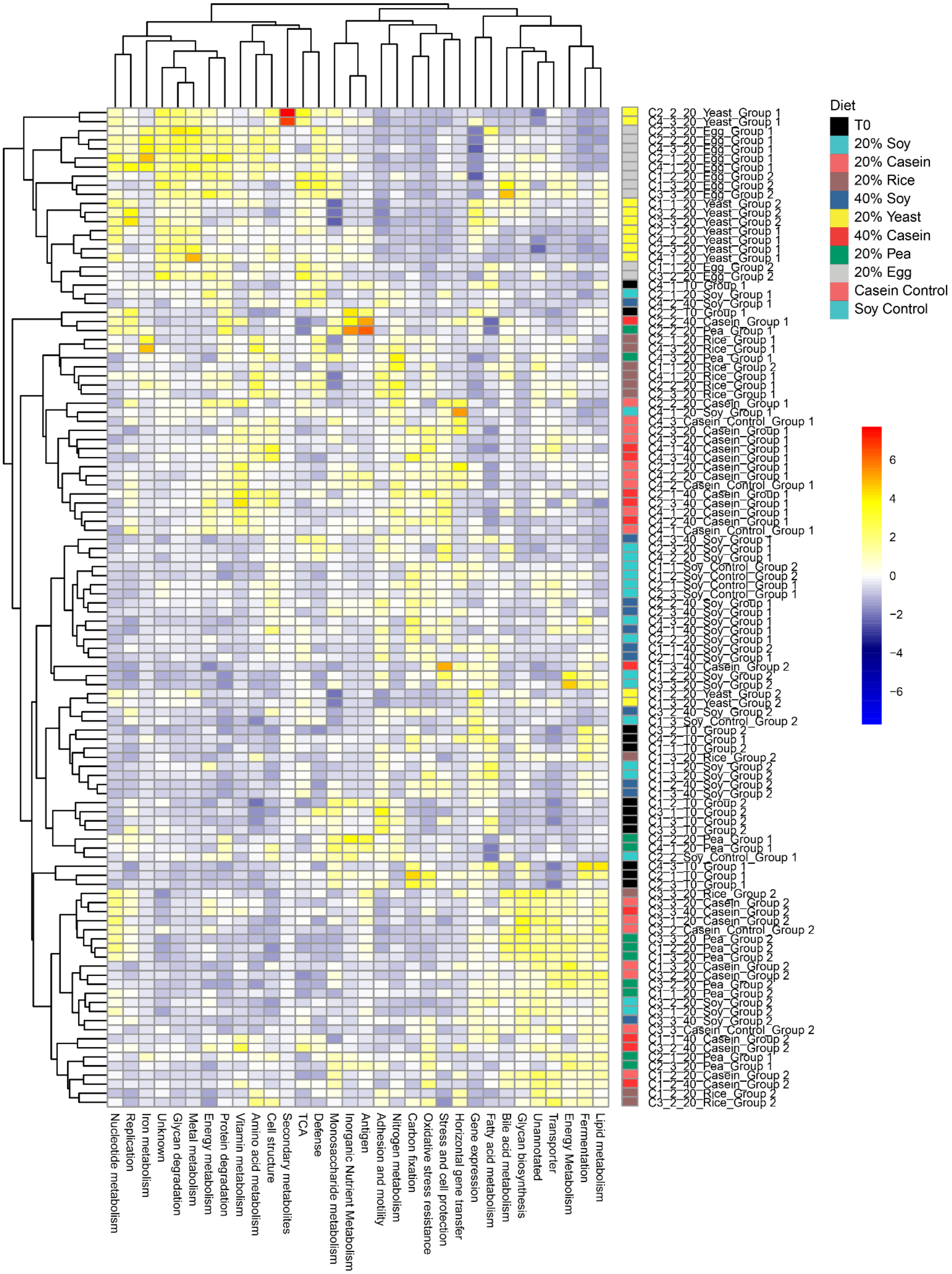


**Supplementary Figure 5: Hierarchical clustering of broad functional categories.** Tabulated abundances of broad functional categories (Extended Data Table 6) were z-scored by feature and then clustered using the ward.D2 method using the pheatmap package in R.

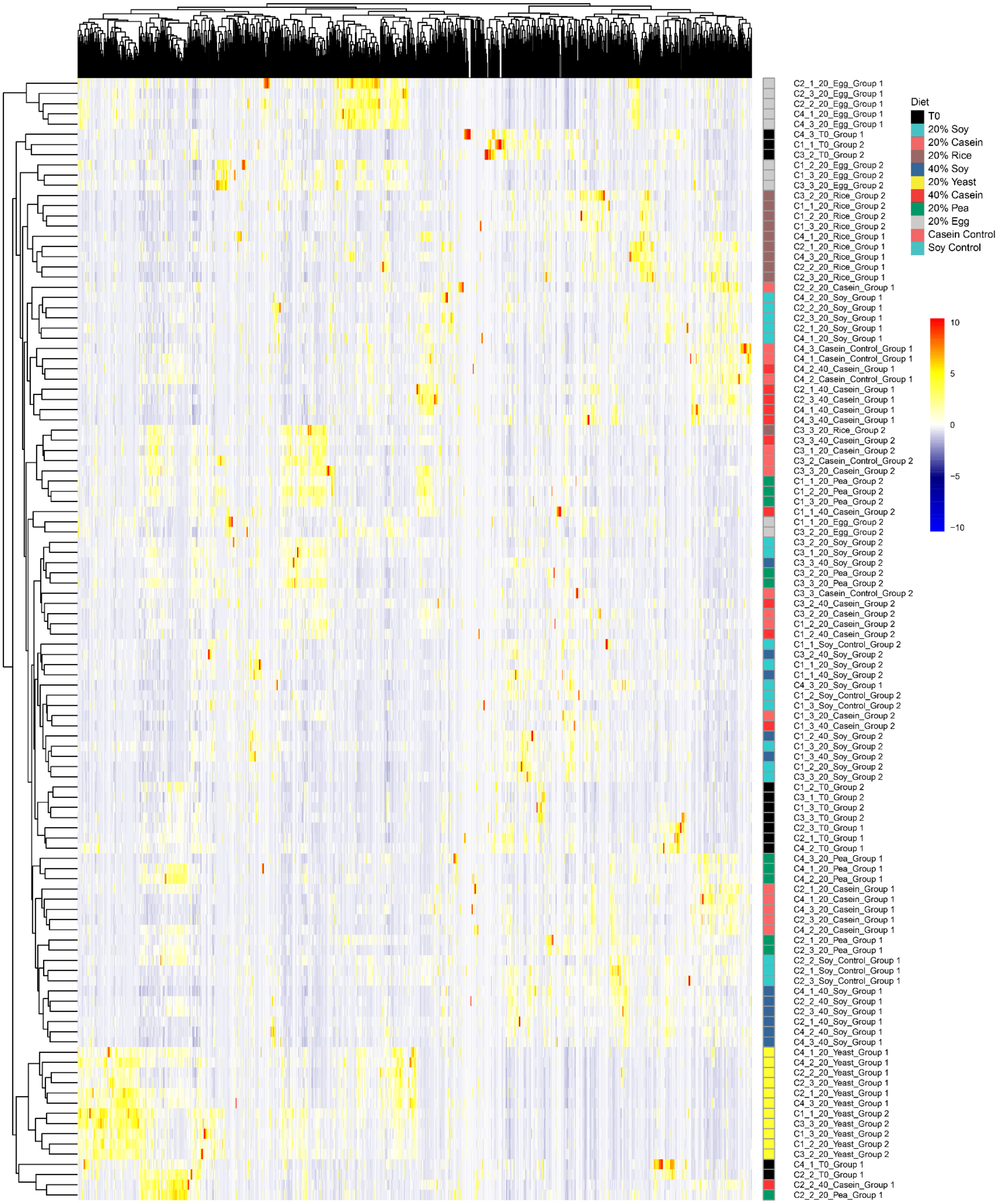


**Supplementary Figure 6: Hierarchical clustering of consensus enzyme names.** Tabulated abundances of consensus enzyme names (Extended Data Table 6) were z-scored by feature and then clustered using the ward.D2 method using the pheatmap package in R.

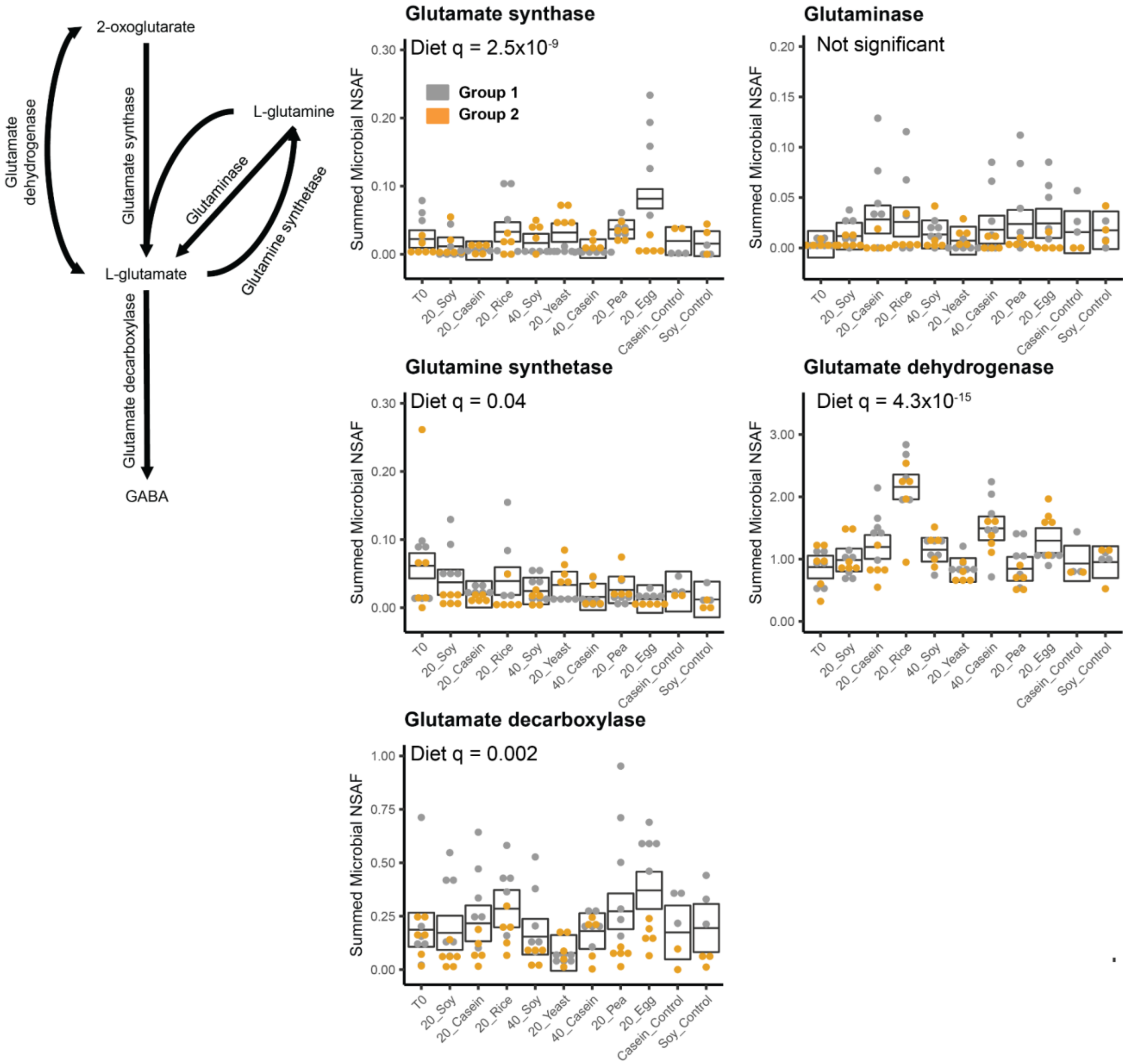


**Supplementary Figure 7: Changes in glutamate and glutamine metabolism due to source of dietary protein:** A reconstruction of the pathways involved in glutamate and glutamine metabolism based on enzymes detected in our metaproteomes. Box plots represent the aggregate abundance of the specific enzymes involved in the pathway. The boxes represent the 95% confidence interval of the mean (line) for each diet from a complete mixed effects model, and the q-values represent the FDR controlled p-values for the diet factor from an ANOVA on these models (q<0.05 indicates significance)(Extended Data Tables 11-12).

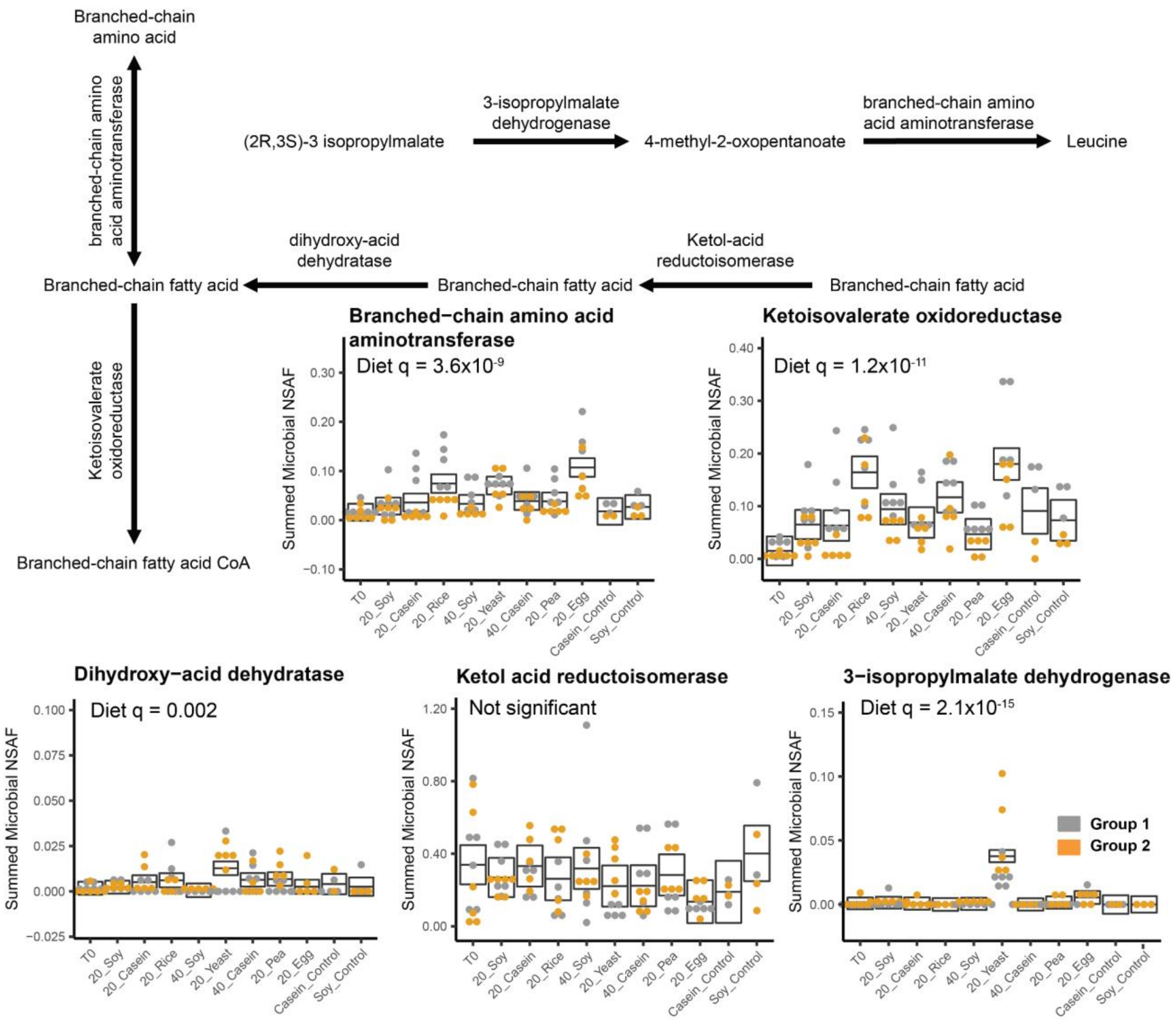


**Supplementary Figure 8: Changes in branched-chain amino acid metabolism due to source of dietary protein.** A reconstruction of the pathways involved in valine, leucine, and isoleucine metabolism based on enzymes detected in our metaproteomes. With the exception of leucine synthesis enzymes, the same enzymes act on all three of these amino acids so we did not try to distinguish them. Box plots represent the aggregate abundance of the specific enzymes involved in the pathway. The boxes represent the 95% confidence interval of the mean (line) for each diet from a complete mixed effects model, and the q-values represent the FDR controlled p-values for the diet factor from an ANOVA on these models ($q<0.05$ indicates significance)(Extended Data Tables 11-12).

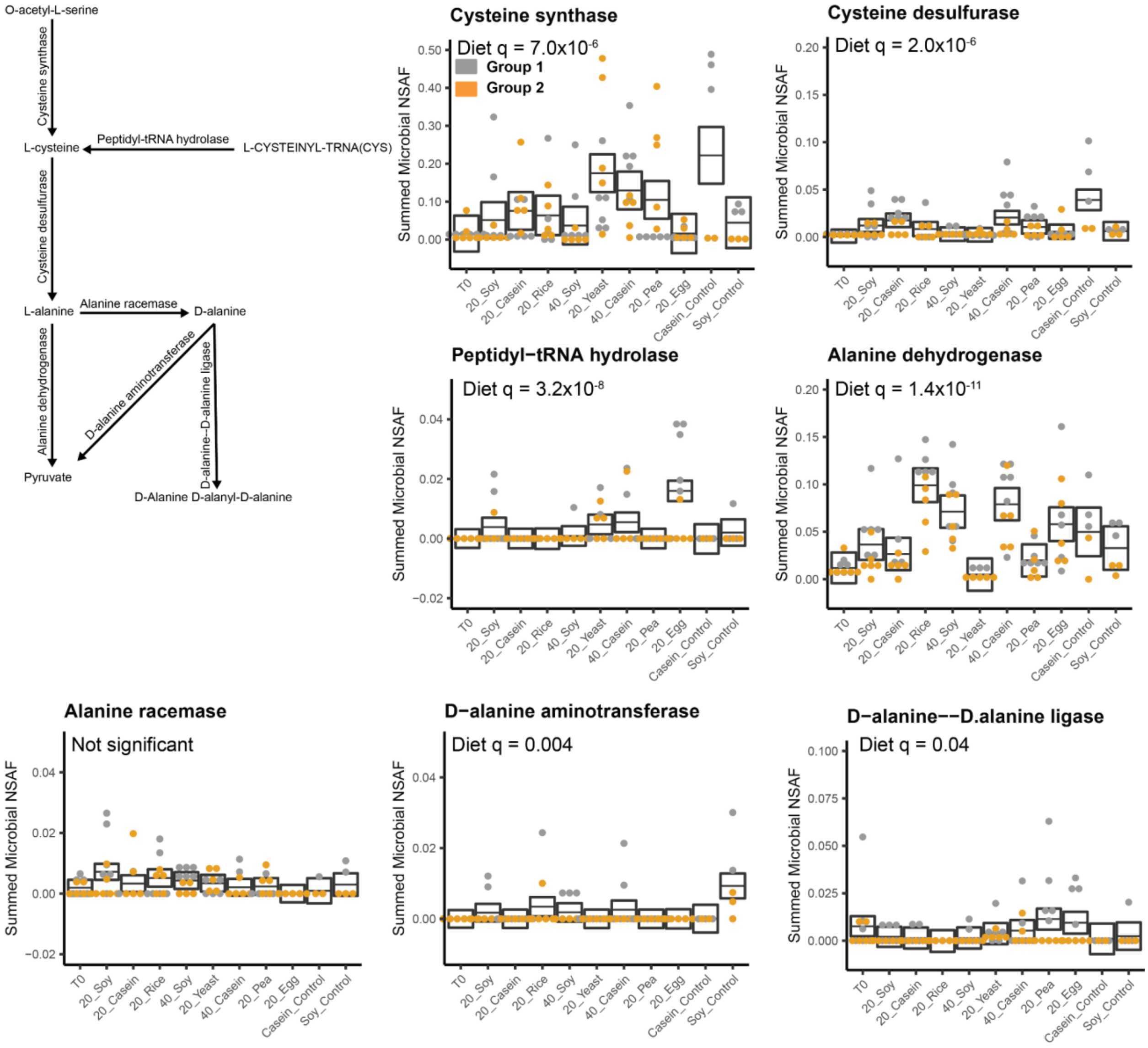

**Supplementary Figure 9: Changes in cysteine and alanine metabolism due to source of dietary protein:** A reconstruction of the pathways involved in cysteine and alanine metabolism based on enzymes detected in our metaproteomes. Box plots represent the aggregate abundance of the specific enzymes involved in the pathway. The boxes represent the 95% confidence interval of the mean (line) for each diet from a complete mixed effects model, and the q-values represent the FDR controlled p-values for the diet factor from an ANOVA on these models ($q<0.05$ indicates significance)(Extended Data Tables 11-12).

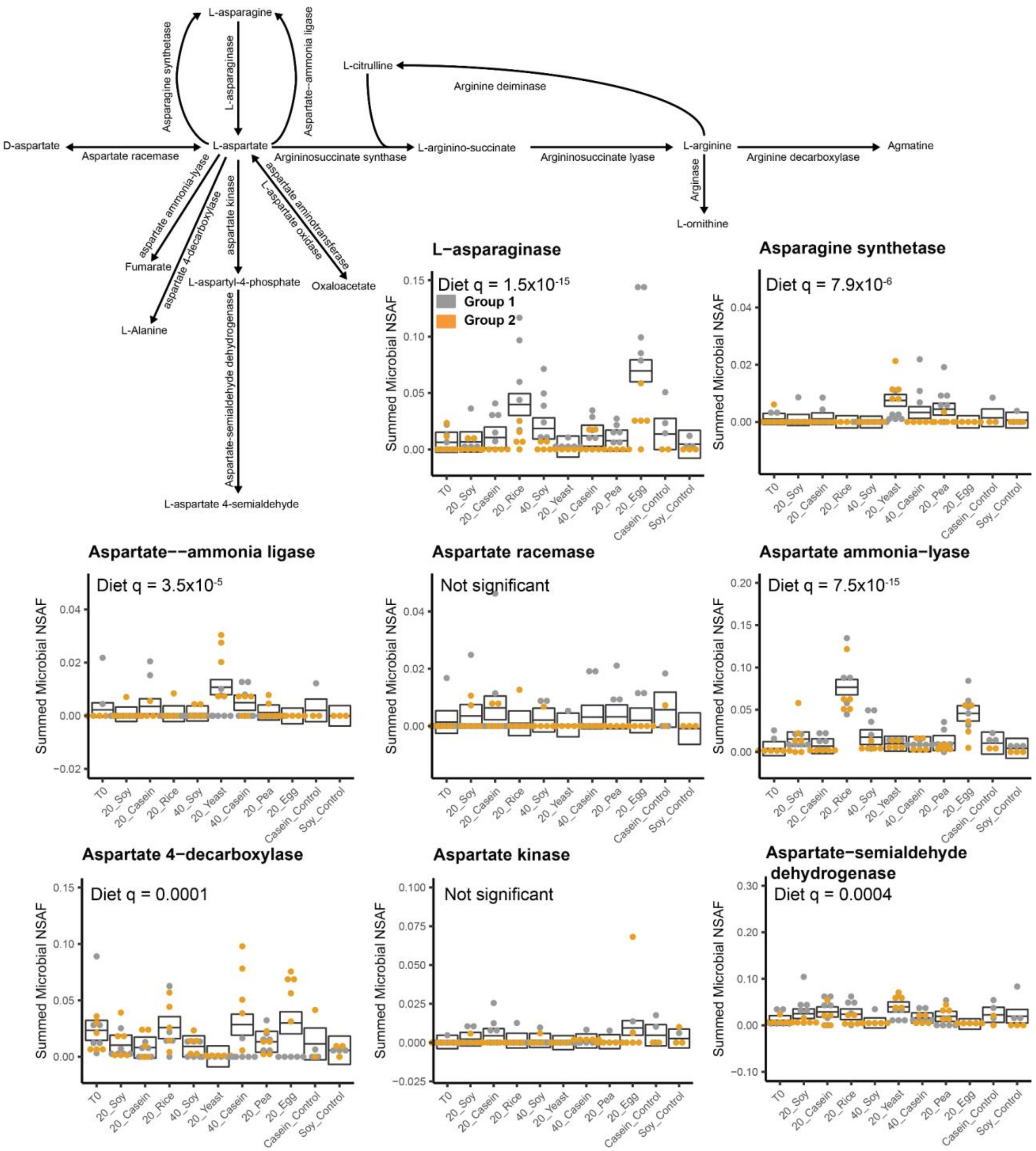

**Supplementary Figure 10: Changes in asparagine, aspartate, and arginine metabolism due to source of dietary protein part 1:**

A reconstruction of the pathways involved in asparagine, aspartate, and arginine metabolism based on enzymes detected in our metaproteomes. Box plots represent the aggregate abundance of the specific enzymes involved in the pathway. The boxes represent the 95% confidence interval of the mean (line) for each diet from a complete mixed effects model, and the q-values represent the FDR controlled p-values for the diet factor from an ANOVA on these models (q<0.05 indicates significance)(Extended Data Tables 11-12).

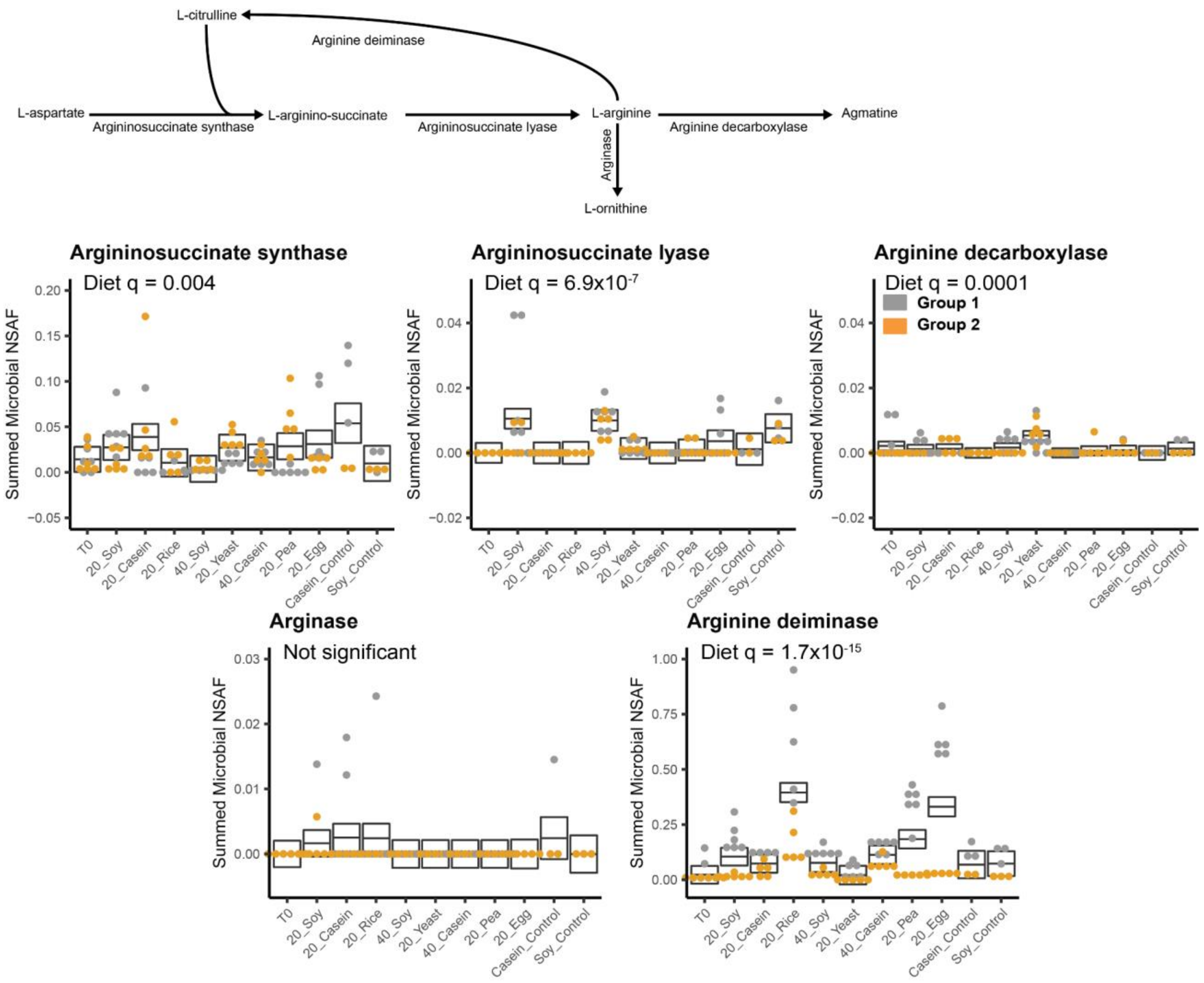


**Supplementary Figure 11: Changes in asparagine, aspartate, and arginine metabolism due to source of dietary protein part 2:**
A reconstruction of the pathways involved in asparagine, aspartate, and arginine metabolism based on enzymes detected in our metaproteomes. Box plots represent the aggregate abundance of the specific enzymes involved in the pathway. The boxes represent the 95% confidence interval of the mean (line) for each diet from a complete mixed effects model, and the q-values represent the FDR controlled p-values for the diet factor from an ANOVA on these models (q<0.05 indicates significance)(Extended Data Tables 11-12).

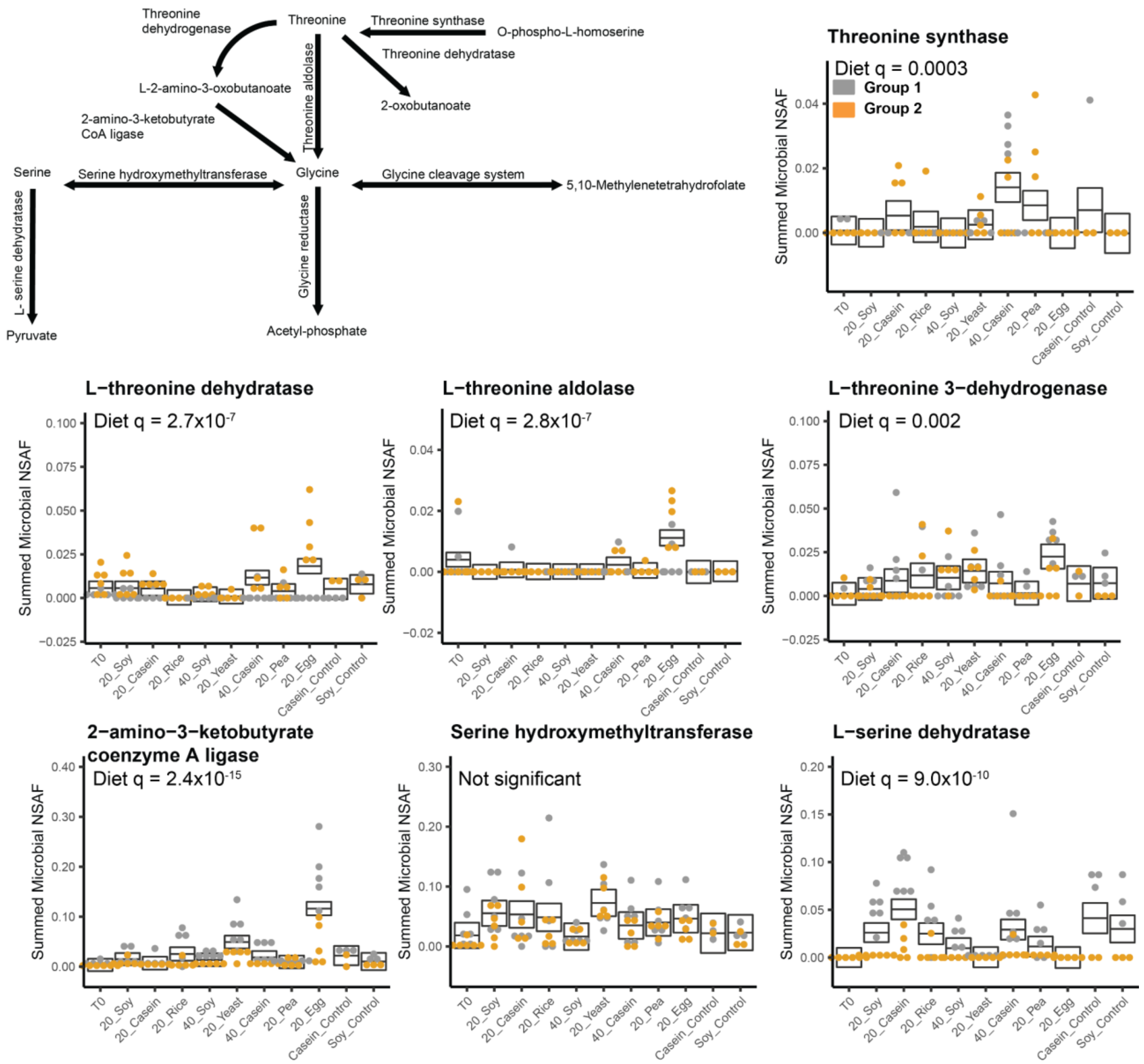


**Supplementary Figure 12: Changes in threonine, glycine, and serine metabolism due to source of dietary protein part 1:**

A reconstruction of the pathways involved in threonine, glycine, and serine metabolism based on enzymes detected in our metaproteomes. Box plots represent the aggregate abundance of the specific enzymes involved in the pathway. The boxes represent the 95% confidence interval of the mean (line) for each diet from a complete mixed effects model, and the q-values represent the FDR controlled p-values for the diet factor from an ANOVA on these models ($q<0.05$ indicates significance)(Extended Data Tables 11-12).

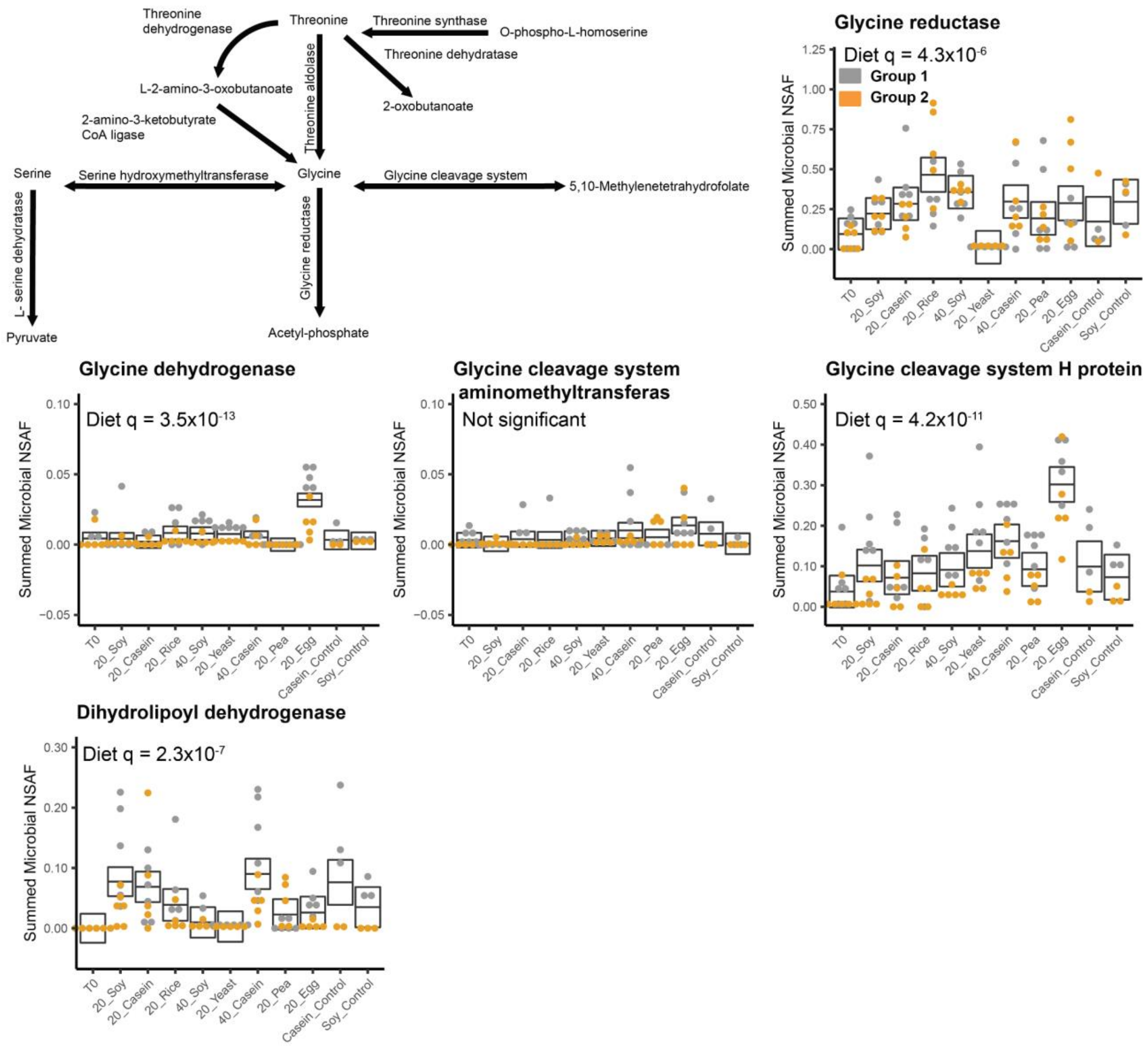


**Supplementary Figure 13: Changes in threonine, glycine, and serine metabolism due to source of dietary protein part 2:**

A reconstruction of the pathways involved in threonine, glycine, and serine metabolism based on enzymes detected in our metaproteomes. Box plots represent the aggregate abundance of the specific enzymes involved in the pathway. The boxes represent the 95% confidence interval of the mean (line) for each diet from a complete mixed effects model, and the q-values represent the FDR controlled p-values for the diet factor from an ANOVA on these models (q<0.05 indicates significance)(Extended Data Tables 11-12).

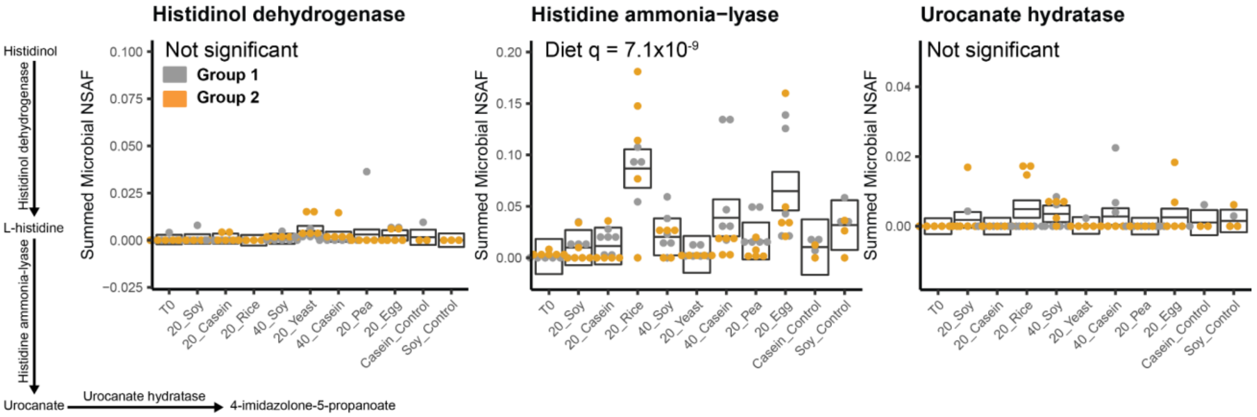


**Supplementary Figure 14: Changes in histidine metabolism due to source of dietary protein:**
A reconstruction of the pathways involved in histidine metabolism based on enzymes detected in our metaproteomes. Box plots represent the aggregate abundance of the specific enzymes involved in the pathway. The boxes represent the 95% confidence interval of the mean (line) for each diet from a complete mixed effects model, and the q-values represent the FDR controlled p-values for the diet factor from an ANOVA on these models (q<0.05 indicates significance)(Extended Data Tables 11-12).

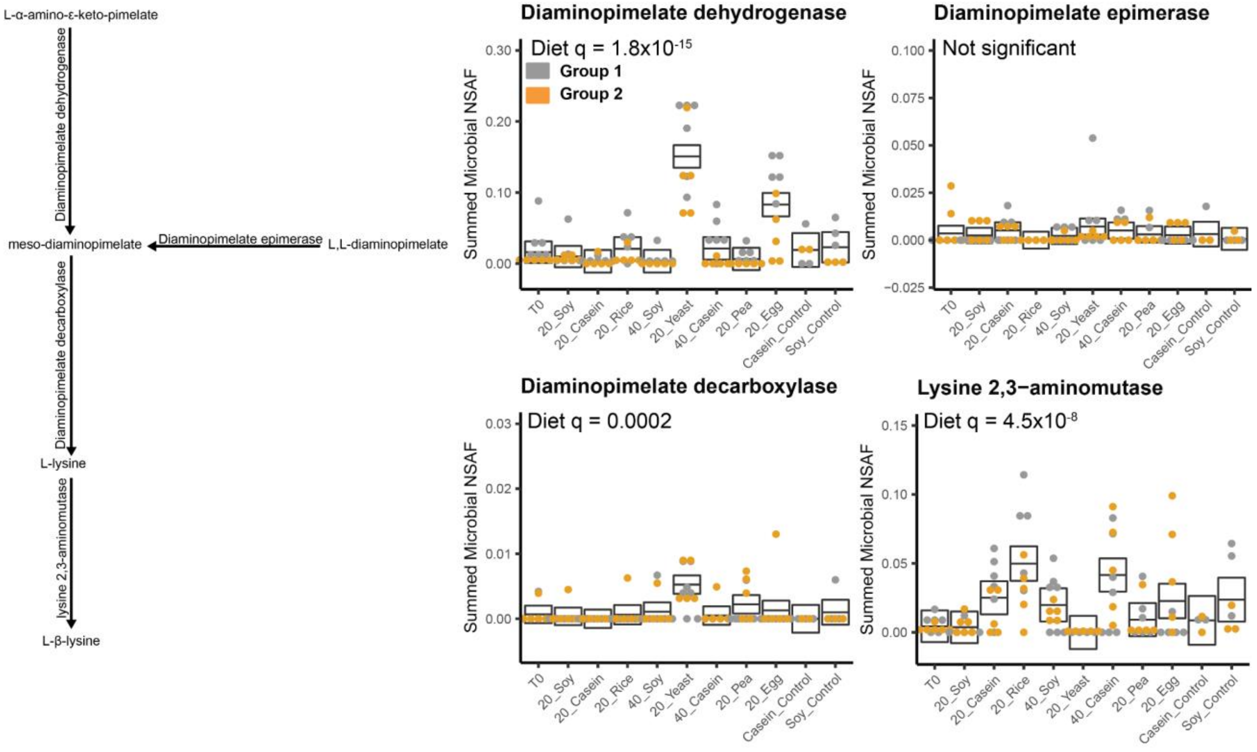


**Supplementary Figure 15: Changes in lysine metabolism due to source of dietary protein:**
A reconstruction of the pathways involved in lysine metabolism based on enzymes detected in our metaproteomes. Box plots represent the aggregate abundance of the specific enzymes involved in the pathway. The boxes represent the 95% confidence interval of the mean (line) for each diet from a complete mixed effects model, and the q-values represent the FDR controlled p-values for the diet factor from an ANOVA on these models (q<0.05 indicates significance)(Extended Data Tables 11-12).

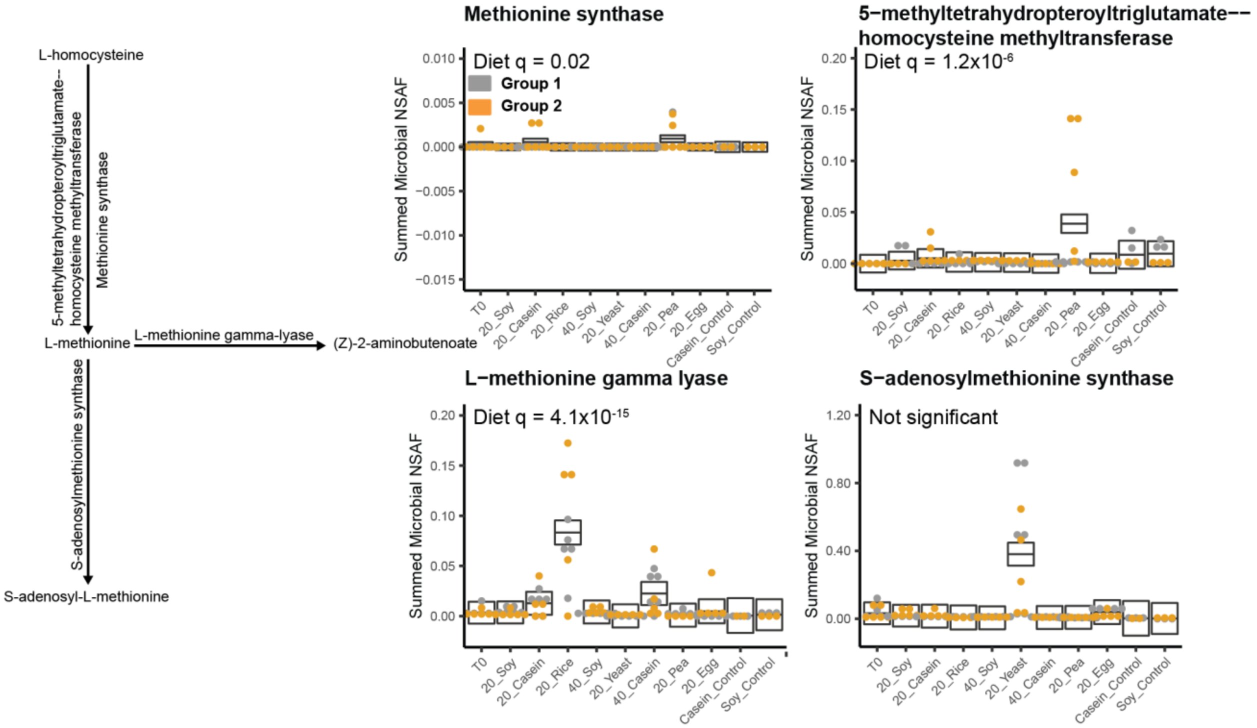


**Supplementary Figure 16: Changes in methionine metabolism due to source of dietary protein:**
A reconstruction of the pathways involved in methionine metabolism based on enzymes detected in our metaproteomes. Box plots represent the aggregate abundance of the specific enzymes involved in the pathway. The boxes represent the 95% confidence interval of the mean (line) for each diet from a complete mixed effects model, and the q-values represent the FDR controlled p-values for the diet factor from an ANOVA on these models ($q<0.05$ indicates significance)(Extended Data Tables 11-12).

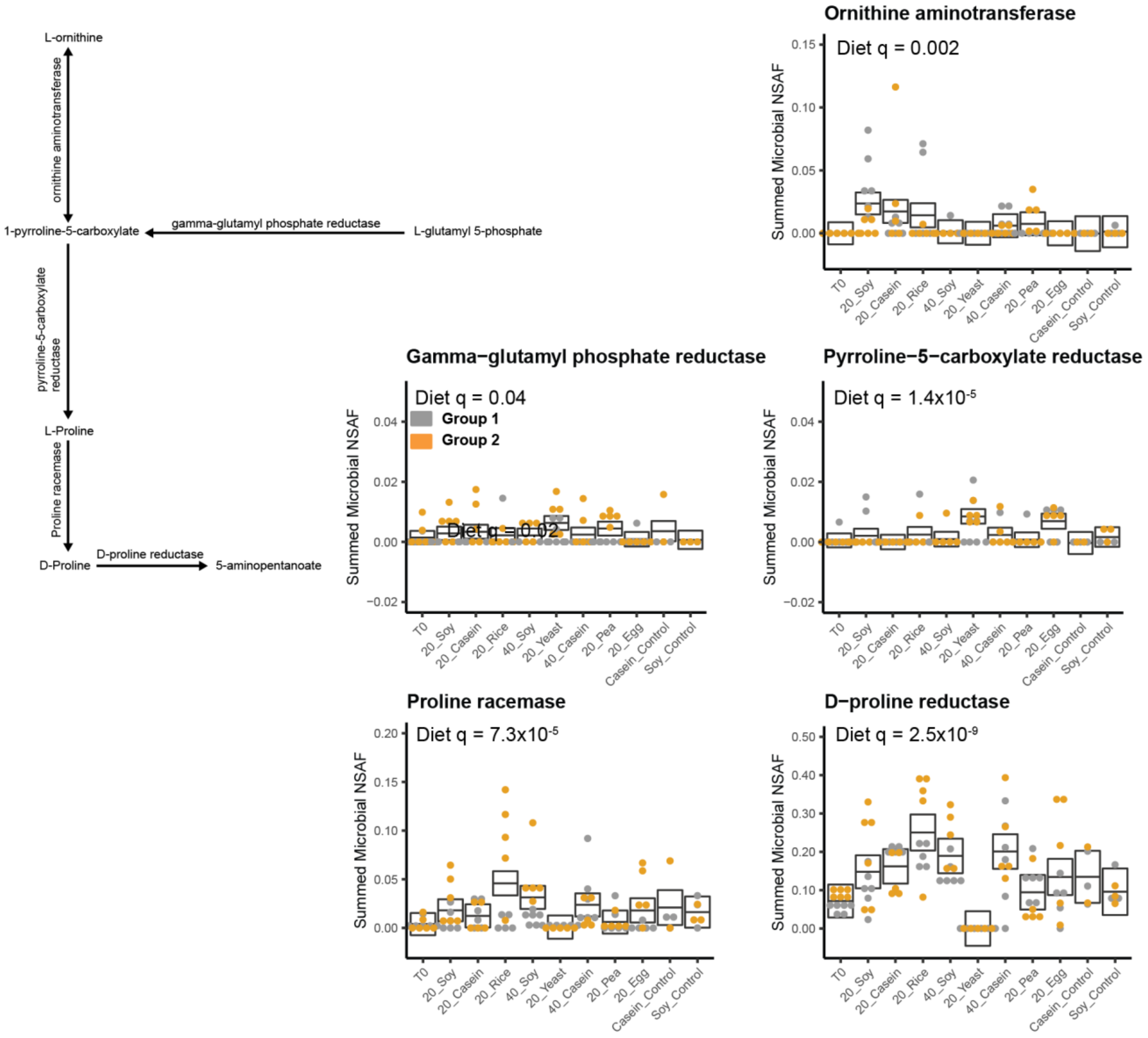


**Supplementary Figure 17: Changes in proline metabolism due to sources of dietary protein:** A reconstruction of the pathways involved in proline metabolism based on enzymes detected in our metaproteomes. Box plots represent the aggregate abundance of the specific enzymes involved in the pathway. The boxes represent the 95% confidence interval of the mean (line) for each diet from a complete mixed effects model, and the q-values represent the FDR controlled p-values for the diet factor from an ANOVA on these models (q<0.05 indicates significance)(Extended Data Tables 11-12).

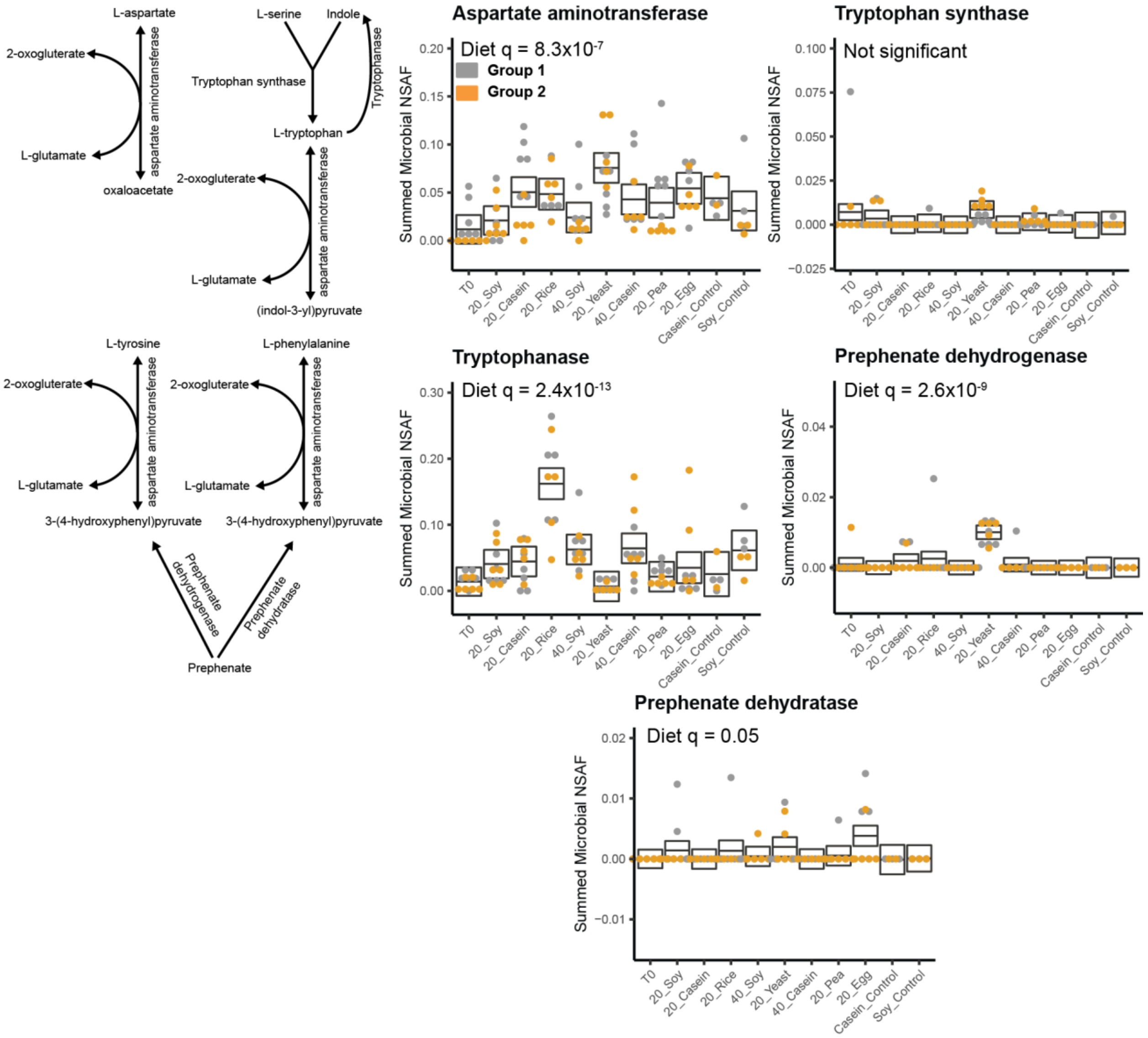


**Supplementary Figure 18: Changes in aspartate, serine, tryptophan, tyrosine, and phenylalanine metabolism due to source of dietary protein:**

A reconstruction of the pathways involved in aspartate, serine, tryptophan, tyrosine, and phenylalanine metabolism based on enzymes detected in our metaproteomes. Box plots represent the aggregate abundance of the specific enzymes involved in the pathway. The boxes represent the 95% confidence interval of the mean (line) for each diet from a complete mixed effects model, and the q-values represent the FDR controlled p-values for the diet factor from an ANOVA on these models ($q<0.05$ indicates significance) (Extended Data Tables 11-12).

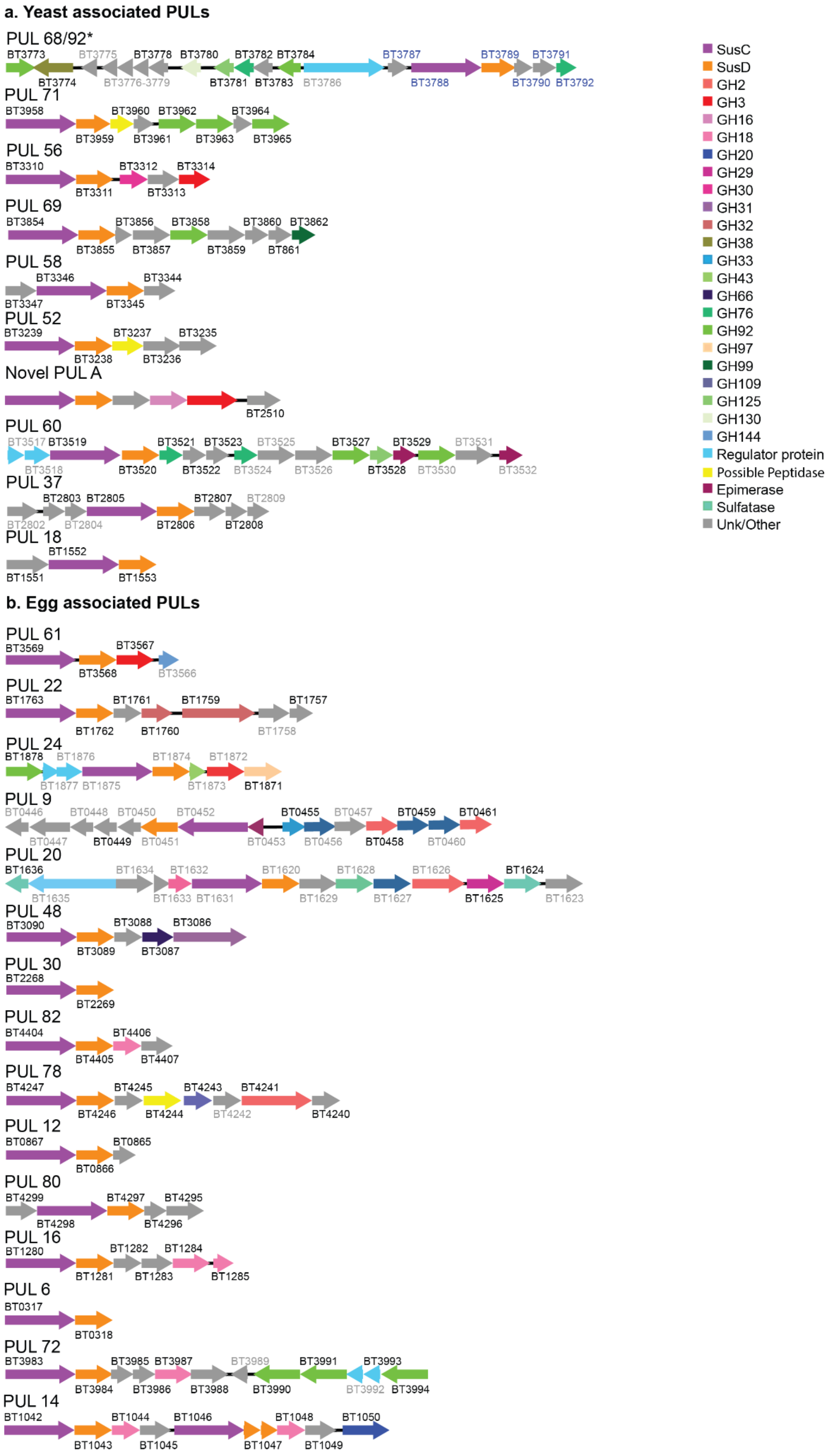
a. Yeast associated PULs
PUL 68/92*
BT3773 BT3775 BT3778 BT3780 BT3782 BT3784 BT3787 BT3789 BT3791
BT3774 BT3776-3779 BT3781 BT3783 BT3786 BT3788 BT3790 BT3792
PUL 71
BT3958 BT3960 BT3962 BT3964
BT3959 BT3961 BT3963 BT3965
PUL 56
BT3310 BT3312 BT3314
BT3311 BT3313
PUL 69
BT3854 BT3856 BT3858 BT3860 BT3862
BT3855 BT3857 BT3859 BT861
PUL 58
BT3346 BT3344
BT3347 BT3345
PUL 52
BT3239 BT3237 BT3235
BT3238 BT3236
Novel PUL A
BT2510
PUL 60
BT3517 BT3519 BT3521 BT3523 BT3525 BT3527 BT3529 BT3531
BT3518 BT3520 BT3522 BT3524 BT3526 BT3528 BT3530 BT3532
PUL 37
BT2803 BT2805 BT2807 BT2809
BT2802 BT2804 BT2806 BT2808
PUL 18
BT1552
BT1551 BT1553
b. Egg associated PULs
PUL 61
BT3569 BT3567
BT3568 BT3566
PUL 22
BT1763 BT1761 BT1759 BT1757
BT1762 BT1760 BT1758
PUL 24
BT1878 BT1876 BT1874 BT1872
BT1877 BT1875 BT1873 BT1871
PUL 9
BT0446 BT0448 BT0450 BT0452 BT0455 BT0457 BT0459 BT0461
BT0447 BT0449 BT0451 BT0453 BT0456 BT0458 BT0460
PUL 20
BT1636 BT1634 BT1632 BT1620 BT1628 BT1626 BT1624
BT1635 BT1633 BT1631 BT1629 BT1627 BT1625 BT1623
PUL 48
BT3090 BT3088 BT3086
BT3089 BT3087
PUL 30
BT2268
BT2269
PUL 82
BT4404 BT4406
BT4405 BT4407
PUL 78
BT4247 BT4245 BT4243 BT4241
BT4246 BT4244 BT4242 BT4240
PUL 12
BT0867 BT0865
BT0866
PUL 80
BT4299 BT4297 BT4295
BT4298 BT4296
PUL 16
BT1280 BT1282 BT1284
BT1281 BT1283 BT1285
PUL 6
BT0317
BT0318
PUL 72
BT3983 BT3985 BT3987 BT3989 BT3991 BT3993
BT3984 BT3986 BT3988 BT3990 BT3992 BT3994
PUL 14
BT1042 BT1044 BT1046 BT1048 BT1050
BT1043 BT1045 BT1047 BT1049
SusC
SusD
GH2
GH3
GH16
GH18
GH20
GH29
GH30
GH31
GH32
GH38
GH33
GH43
GH66
GH76
GH92
GH97
GH99
GH109
GH125
GH130
GH144
Regulator protein
Possible Peptidase
Epimerase
Sulfatase
Unk/Other

**Supplementary Figure 19: Distinct polysaccharide utilization loci (PULs) are expressed by *B. theta* between the egg white and yeast diet.** Graphical representation of PUL gene neighborhoods detected in the metaproteome. PUL identifiers are the literature derived PUL identifiers from PULDB[8]**,** but the PUL structure was verified in RAST using the *B. theta* genome assembled from our metagenome. Metagenome identifiers were cross referenced to BT numbers from previous PUL papers. If the BT number is black, it is detected in the metaproteome, if gray it is not detected in our metaproteome but detected in our genome, and if blue it means that we did not have those genes in our genome but instead detected homologs with the exact same gene neighborhood structure and similar gene percent identity. (a) PUL operons detected in our metaproteome that were increased in the yeast diet relative to the egg white diet.(c) PUL operons detected in our metaproteome that were increased in the egg white diet relative to the yeast diet.

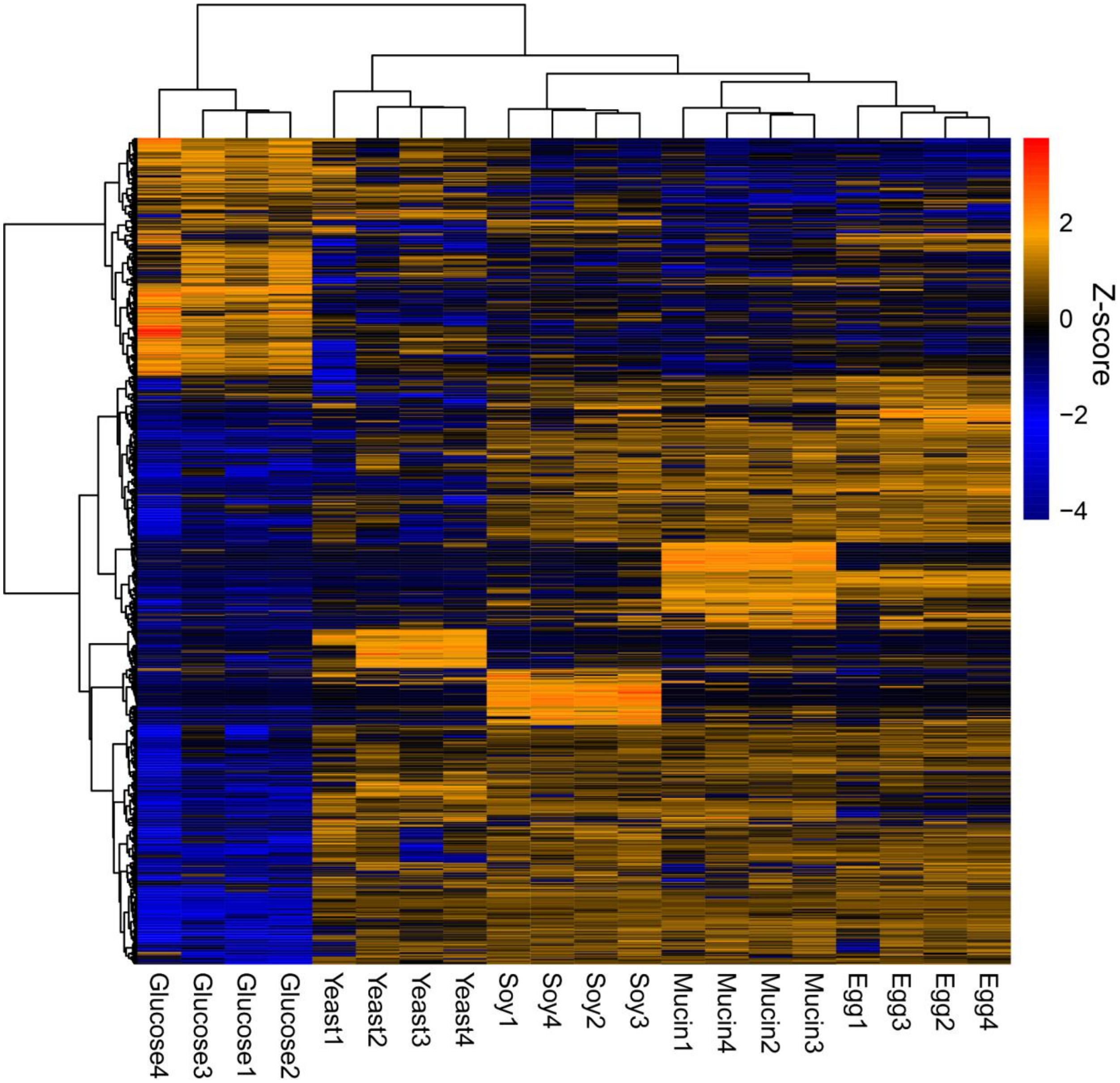


**Supplementary Figure 20: Significantly different proteins in *in vitro* proteomes of *B. theta* clustered by growth medium.** Clustered heatmap of the *in vitro* proteomes of *B. theta* after z-score standardization and removal of non-significant proteins after testing by ANOVA (q<0.05). We generated dendrograms using the ward clustering algorithm on euclidean distances.
